# Sex differences in inter-individual gene expression variability across human tissues

**DOI:** 10.1101/2022.08.09.503366

**Authors:** Samuel Khodursky, Caroline S. Jiang, Eric B. Zheng, Roger Vaughan, Daniel R. Schrider, Li Zhao

## Abstract

Understanding phenotypic sex differences has long been a goal of biology from both a medical and evolutionary perspective. Although much attention has been paid to mean differences in phenotype between the sexes, little is known about sex differences in phenotypic variability. To gain insight into sex differences in inter-individual variability at the molecular level, we analyzed RNA-seq data from 43 tissues from the Genotype-Tissue Expression project (GTEx). Within each tissue, we identified genes that show sex differences in gene expression variability. We found that these sex-differentially variable (SDV) genes are associated with various important biological functions, including sex hormone response, immune response, and other signaling pathways. By analyzing single-cell RNA sequencing data collected from breast epithelial cells, we found that genes with sex differences in gene expression variability in breast tissue tend to be expressed in a cell-type-specific manner. We looked for an association between SDV expression and Graves’ disease, a well-known heavily female-biased disease, and found a significant enrichment of Graves’ associated genes among genes with higher variability in females in thyroid tissue. This suggests a possible role for SDV expression in the context of sex-biased disease. We then examined the evolutionary constraints acting on genes with sex differences in variability and found that they exhibit evidence of increased selective constraint. Through analysis of sex-biased eQTL data, we found evidence that SDV expression may have a genetic basis. Finally, we propose a simple evolutionary model for the emergence of sex-differentially variable expression from sex-specific constraints.

## Introduction

Identifying, characterizing, and understanding phenotypic differences between the sexes has long been a major goal of biology. A very wide variety of traits in humans show significant sex differences including morphological, behavioral, metabolic, immune, and disease traits (Chella Krishnan et al., 2018; Choleris et al., 2018; Klein and Flanagan, 2016; Ober et al., 2008). These trait differences are likely in part due to sex differences in gene expression and regulation (Naqvi et al., 2019; Parsch and Ellegren, 2013; Williams and Carroll, 2009). As a phenotype, gene expression is both relatively straightforward to measure and highly quantitative. As a result, many studies have examined differences in mean gene expression levels between the sexes in many species (Catalán et al., 2012; Gershoni and Pietrokovski, 2017; Khodursky et al., 2020; Lopes-Ramos et al., 2020; Naqvi et al., 2019; Oliva et al., 2020; Parsch and Ellegren, 2013). Sex-biased expression (SBE) is thought to arise from sex-specific genetic architectures which are ultimately established by sex-determination systems initiated during early development (Williams and Carroll, 2009). In addition to underlying sexually dimorphic phenotypes, genes with SBE have been found to possess many interesting evolutionary and genomic properties (Parsch and Ellegren 2013). For example, SBE has been associated with increased rates of evolution at both the coding and expression levels in *Drosophila* and other animals (Khodursky et al., 2020; Meisel et al., 2012; Parsch and Ellegren, 2013; Ranz et al., 2003). In humans, genes with SBE have been associated with reduced selective constraint (Gershoni and Pietrokovski, 2017; Naqvi et al., 2019). Additionally, genes with SBE have been shown to be distributed across the genome in a non-random fashion with respect to the X chromosome – thereby linking the evolution of sex-biased genes to the unique evolutionary forces experienced by the X chromosome (Khodursky et al., 2020; Oliva et al., 2020; Parisi et al., 2003).

Relative to sex differences in phenotypic means, sex differences in phenotypic variabilities have been less thoroughly studied. In the biomedical sciences there has been a long-standing assumption that females must generally be more phenotypically variable due to the menstrual or estrous cycles; however, there has been little empirical evidence to support this assumption (Beery and Zucker, 2011). In fact, a meta-analysis of microarray studies in mice and humans revealed that males may have slightly higher expression variability (Itoh and Arnold, 2015) and a meta-analysis of various traits in mice indicated that sex differences in phenotypic variability are trait dependent (Zajitschek et al., 2020). Despite growing interest in understanding sex differences in phenotypic variability, the biological and evolutionary properties of genes with sex differences in gene expression variability have not, to our knowledge, been explored in humans. Here, we define genes with sex-differentially variable (SDV) expression to be genes which show significant differences in inter-individual gene expression variability between the sexes. From a medical standpoint, genes with SDV expression may underlie diseases for which there are sex differences in prevalence or outcome. For example, one could imagine a disease partially caused by the aberrant expression of a certain gene (akin to the liability threshold model; (Falconer, 1967)). If one sex has more variable expression of the gene in a normal healthy population, this may help explain a higher prevalence of that disease in that sex. Additionally, since more variable expression has been associated with reduced effectiveness of drug targets it is possible that SDV expression may account for sex differences in drug effectiveness, which are known to exist (Simonovsky et al., 2019; Soldin and Mattison, 2009).

Understanding the evolutionary properties of these genes may give us novel insight into the evolution of sex differences. For example, genes with sex differences in expression variability may evolve under sex-specific constraints (e.g., purifying or stabilizing selection) and be of particular functional importance for a given sex. Here we examine the biological properties and evolutionary constraints associated with SDV expression across 43 human tissues.

## Results

### Identification and characterization of genes with sex differences in gene expression variability

We obtained RNA-seq count data for 43 tissues from the GTEx project (version 8). Since we sought to identify genes with sex differences in gene expression variability, we first compared several statistical methods using power simulations. Specifically, we compared three tests: the Fligner-Killeen test, Levene’s test, and a test based on negative binomial generalized additive models for location, scale and shape (GAMLSS) (Rigby and Stasinopoulos, 2005). Since RNA-seq count data is known to follow an approximately negative binomial distribution (Robinson and Smyth, 2007; Robinson et al., 2010), we performed power simulations by drawing random samples from negative binomial distributions with identical means but different variances by varying the overdispersion term (see methods). At each parameter combination, 1000 pairs of samples were drawn, and tests were performed at an alpha of 0.05. We found that GAMLSS had the highest power across a range of parameters, and this was especially true when testing for differences in variance between samples of unequal size (**Supplementary Figure 1A-D**). Both GAMLSS and Levene’s test managed to maintain an adequate false positive rate (FPR) across a range of parameters. Beyond increased power, GAMLSS offers other advantages over the other two tests: GAMLSS negative binomial models allow for the independent parametrization of both the mean and the non-Poisson variation (overdispersion) terms as a function of variables and covariates of interest, including library size (de Jong et al., 2019; Rigby and Stasinopoulos, 2005).

When considering the inter-sample variation of gene expression, it is natural to consider the coefficient of variation: the standard deviation divided by the mean. This provides a mean-normalized statistic for variability. Notably, it is thought that the Poisson component of the coefficient of variation in RNA-seq data is due to technical variation while the overdispersion component of the coefficient of variation contains the true biological variation (McCarthy et al., 2012). The overdispersion parameter fit by GAMLSS can therefore be considered to be the square of the biological coefficient of variation (de Jong et al., 2019; McCarthy et al., 2012). Observed overdispersion values and effect sizes in GTEx data were similar to the simulated values, as determined by models fit using GAMLSS (**Supplementary Figure 1E**).

To determine the effect of sex on gene expression variation we fit three negative binomial generalized additive models using the R library ‘gamlss’ for each expressed gene in each tissue (Rigby and Stasinopoulos, 2005). To reduce the risk of confounding due to unequal sampling across various populations, we limited our analysis to European-American individuals. Significance of the effect of sex on overdispersion and mean expression level was determined through a pair of likelihood-ratio tests (see methods). All models included covariates for age, RNA integrity number (RIN), ischemic time, and the first five genotype principal components (to control for population structure) for both mean and overdispersion terms. We initially considered genes to be sex-differentially variable (SDV) if the effect of sex on overdispersion was significant at FDR<0.05. Since we observed that outliers dramatically influenced estimates of variation in some genes, as a secondary test we performed randomization tests by randomly permuting the sex labels on genes with initially significant coefficients for the effect of sex on overdispersion and refitting the GAMLSS models (see methods) (**Figure 1A**). Coefficients for the effect of sex on overdispersion were saved after each permutation. We used 1000 rounds of permutation to generate empirical null distributions for the effect of sex on overdispersion. Initially significant genes with FDR-adjusted empirical p-values greater than 0.05 in the permutation test were no longer considered to be SDV (see methods). Results from all tested genes are included in **Supplementary Table 1**.

**Figure 1.**
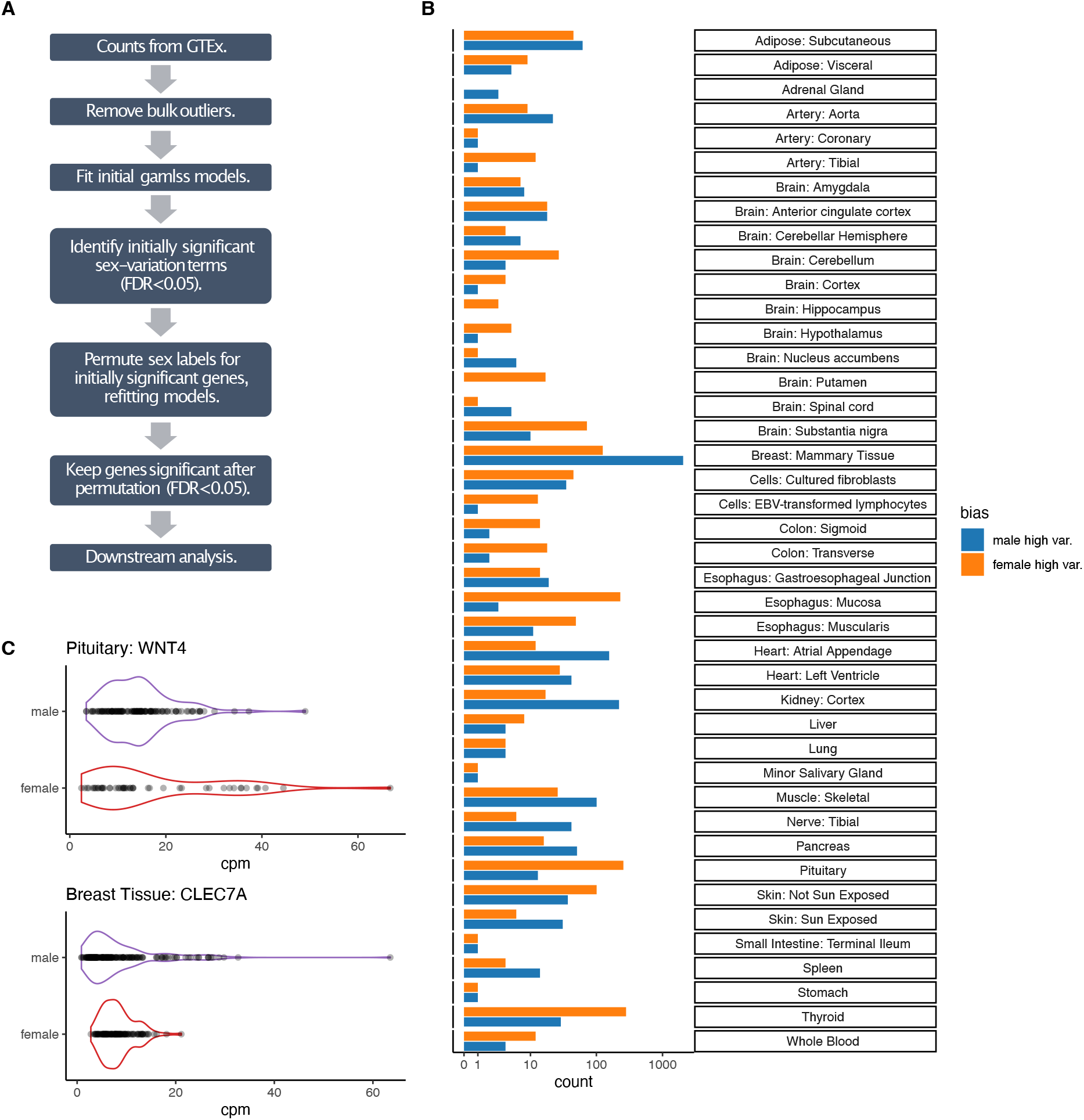
The identification of genes with sex-differentially-variable gene expression. A) Outline of pipeline to identify SDV genes. B) The number of genes with SDV expression identified in each tissue. The X-axis has a pseudo-log scale. C) Two examples of SDV genes with higher variation in females (WNT4 in the pituitary gland) and in males (CLEC7A in breast tissue), respectively.

Across 43 tissues, we identified 3895 unique genes with SDV expression in at least one tissue out of a total 22054 tested genes (**Figure 1A, B and Supplementary Table 1**). We were able to identify SDV genes in 42 out of the 43 tested tissues (**Figure 1B**). Examples of two genes with SDV expression in the pituitary gland and breast tissue are shown in **Figure 1C**. A total of 2792 unique genes showed higher variability in males in at least one tissue, while 1382 genes showed higher variability in females in at least one tissue. This imbalance between the sexes was largely driven by breast tissue. Surprisingly, 279 genes had higher variability in males in one tissue and higher variability in females in at least one other tissue. The vast majority (3347) of SDV genes only showed SDV expression in one tissue while 548 genes showed SDV expression in multiple tissues.

Breast tissue had the largest number of SDV genes with 2049 genes showing significantly higher variation in males and 125 genes showing significantly higher variation in females (**Figure 1B**). Apart from a few tissues, SDV genes were not consistently expressed at higher or lower levels than non-SDV genes across tissues (in terms of both CPM and TPM), suggesting that our discovery of SDV genes was not significantly biased towards genes of high or low expression (**Supplementary Figure 2**).

Given that differences in variation are generally difficult to detect between groups, we performed a replication analysis using data from breast tissue to help validate our statistical approach. First, we randomly split individuals into discovery and replication sets – 70% of males and females were placed in the discovery set while the rest were placed in the replication set. We identified genes with sex differences in overdispersion in the discovery set using the combined likelihood ratio test and permutation approach described above (both at FDR<0.05). These genes were then tested in the replication set: genes with p-values for the overdispersion term less than 0.05 in both the likelihood ratio test and permutation test were considered ‘replicated’ if the sex with higher variability in the discovery set was also more variable in the replication set. We observed a replication rate of 21.9% (**Supplementary Figure 3**). This replication rate is limited by reduced power due to the relatively small size of the replication set. To determine the significance of this replication rate we generated an empirical null distribution under the null hypothesis that our statistical approach was simply randomly sampling genes. Therefore, we randomly sampled genes from the discovery set and calculated the replication rate for each sample. The median replication rate under the null hypothesis was found to be 8.4% and out of 10,000 random samples none had a replication rate higher than 11.3% (**Supplementary Figure 3**). This null replication rate should not be considered as a type 1 error rate. Since we are randomly sampling genes from the set of all expressed genes, we are also capturing some true SDV genes in each sample. One would expect the replication rate under random sampling to be higher than 5% even if the FDR is properly controlled at 5%.

Since our GAMLSS framework allowed us to simultaneously test for sex differences in both mean and overdispersion we examined the relationship between SDV genes and genes with sex-biased expression (SBE). In several tissues a majority of SDV genes also displayed sex-biased expression (**Supplementary Figure 4**). Although a minority of SDV genes in breast tissue, 466 genes with significant sex differences in both mean and overdispersion showed discordance, with higher overdispersion but lower mean expression in one sex versus the other (**Supplementary Figure 4**). Specifically, 387 of those genes simultaneously had significantly higher overdispersion in males and significantly higher mean expression in females.

Given that genes with sex-biased expression had been shown to be overrepresented on the X chromosome in a variety of tissues and species (Khodursky et al., 2020; Oliva et al., 2020; Parisi et al., 2003), we wondered if SDV genes were similarly enriched on the X chromosome. However, we found no consistent enrichment of SDV genes on the X chromosome; the enrichment patterns were highly tissue-dependent (**Supplementary Figure 5**).

To identify biological processes associated with SDV gene expression, we performed an overrepresentation analysis using gene sets obtained from the hallmark gene set collection of MSigDB (Liberzon et al., 2015; Subramanian et al., 2005). The hallmark gene set collection consists of non-redundant and refined gene sets associated with various biological processes and states (Liberzon et al., 2015). The hallmark gene set collection is ideal for our analysis since we identified few SDV genes in most tissues – a larger, more redundant, and less refined collection would introduce more noise while increasing the multiple testing burden for our overrepresentation analysis. We first performed an enrichment analysis at the tissue level and aggregated results across tissues using an empirical approach based on Fisher’s combined probability method (**Figure 2A, B, C**) (see methods). Following the meta-analysis, we found that genes with higher variation in males were overrepresented in sets related to estrogen and androgen response, immune response, and cancer-associated pathways (**Figure 2A**). Genes with increased variation in females were similarly overrepresented in sets related to immune response, estrogen response, and cancer-associated pathways (**Figure 2A)**. Notably, there was overlap in multiple sets between the two sexes even though the lists of SDV genes did not show much overlap. At the single-tissue level, genes with higher variation in males in breast tissue were overrepresented in sets related to estrogen response and immune response (**Figure 2B**). This suggests that genes of particular biological importance to one sex (genes related to estrogen response in female breast tissue) may show higher variation in the opposite sex. To identify transcription factors whose targets are overrepresented in the set of SDV genes we downloaded the GTRD dataset from MSigDB (Liberzon et al., 2015; Subramanian et al., 2005). The GTRD collection contains sets of transcription factors and their predicted binding targets (within the promoter region) (Kolmykov et al., 2021). We first performed an enrichment analysis at the individual tissue level, followed by a meta-analysis across tissues using the same modified Fisher’s combined probability approach mentioned above (see methods). Following the meta-analysis, we found that genes with higher variability in females were significantly enriched for targets of MAML1 and GLI1 (**Figure 2C**). Genes with higher variability in males were significantly enriched for targets of KMT2D. GLI1 is a transcription factor that is part of the Hedgehog signaling pathway, which is involved in differentiation during development. However, in fully differentiated tissues GLI1 can act as an oncogene has been implicated in a wide variety of cancers, including glioblastoma, breast cancer, and colon cancer (Avery et al., 2021; Clement et al., 2007; Sun et al., 2014). MAML1 is a transcription factor involved in Notch signaling and is a transcriptional co-activator of Notch receptors (Wu et al., 2000). Additionally, MAML1 has been shown to interact cooperatively with GLI1 to regulate hedgehog signaling (Quaranta et al., 2017). KMT2D is a broadly expressed histone methyltransferase that is believed to function as a tumor suppressor and is frequently mutated in a wide variety of cancers (Rao and Dou, 2015).

**Figure 2.**
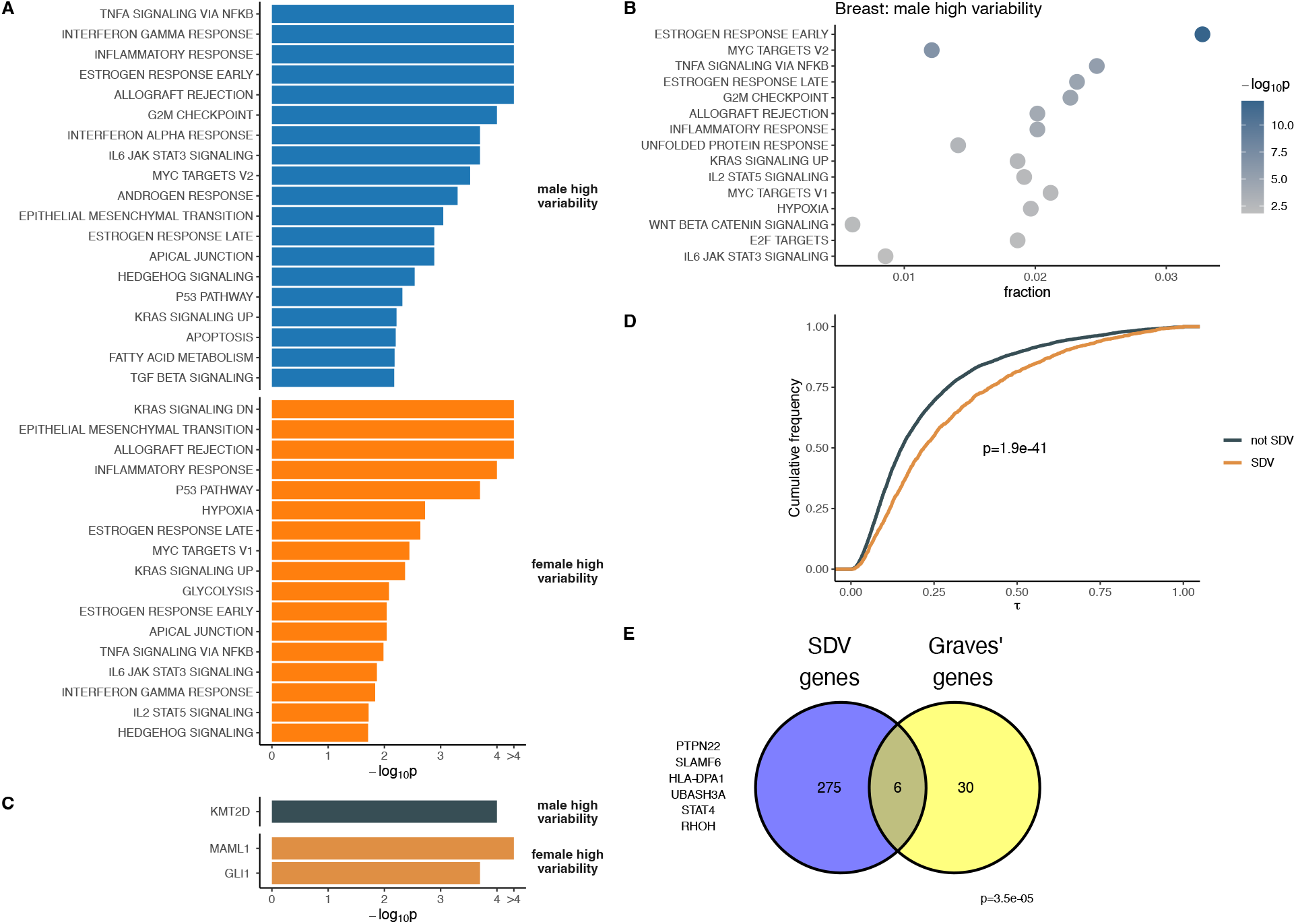
Biological properties of SDV genes. A) Following meta-analysis SDV genes with higher variation in males and females are enriched for terms related to various signaling pathways, immune response, and sex-hormone response. B) Genes with higher variability in males are enriched for terms related to estrogen and immune response in breast tissue. C) Following meta-analysis genes with higher variability in females are enriched for targets of two transcription factors. D) SDV genes identified in breast tissue show highly cell-type specific expression in breast epithelial tissue (p-value calculated using the Mann-Whitney U test). Higher τ indicates greater cell-type specificity. E) Genes with higher variability in females in the thyroid are significantly enriched for genes associated with Graves’ disease (p-value calculated using a hypergeometric test).

### Single-cell RNA-seq data from breast epithelial cells shows that genes with SDV expression tend to be expressed in a cell type-specific manner

To get further biological insight into genes with SDV expression in breast tissue, which had the most SDV genes identified out of any tissue by far, we analyzed a previously published single cell RNA-seq dataset (Nguyen et al., 2018). With count data from individual 4 (female), we applied a standard single cell RNA-seq pipeline using the ‘R’ library ‘Seurat’ (Hao et al., 2020)(see methods). Our analysis recovered the three major cell types found in the breast epithelium: type 1 and type 2 luminal cells along with basal cells (Nguyen et al., 2018)(**Supplementary Figure 6**). To determine the cell type specificity of each gene we combined raw count data from each cell to generate a pseudo-bulk count matrix for each cell type and calculated CPM values for each gene. We then applied a standard measure of tissue specificity, *τ*, to the log-transformed CPM values across the three cell types (see methods).

We found that genes exhibiting SDV expression were expressed in a significantly more cell-type-specific manner than non-SDV genes (p=1.9e-41, Mann-Whitney U test)(**Figure 2D**). If there exist significant sex differences in inter-individual cell-type composition variation, such cell-type-specific expression could lead to sex differences in expression variability in bulk tissue samples. Overall, these results suggest that SDV expression may be partly due to sex differences in inter-individual cell-type composition variation.

### SDV genes in the thyroid with higher variability in females are associated with Graves’ disease

Given the preponderance of sex-biased diseases (Mauvais-Jarvis et al., 2020), we wondered if genes with SDV expression could be associated with a classically sex-biased disease. Hence, we decided to examine Graves’ disease: a common autoimmune disease of the thyroid which results in hyperthyroidism (McIver and Morris, 1998). The incidence of Graves’ disease is almost ten-fold higher in females than in males (McIver and Morris, 1998). We obtained GWAS summary data from the NHGRI-EBI GWAS catalog compiled from 5 studies on the genetics of Graves’ disease in various populations (Buniello et al., 2019; Chu et al., 2011; Cooper et al., 2012; Nakabayashi et al., 2011; Sakaue et al., 2021; Zhao et al., 2013). Of the 36 genes implicated in Graves’ disease which were expressed in thyroid tissue, 6 displayed SDV expression with higher variation in females. This corresponds to a significant 9-fold enrichment of Graves’ associated genes among SDV genes with higher variability in females (p=3.5e-5, hypergeometric test) (Figure 2E). Although many of the genes implicated in Graves’ disease are expressed in immune cells, a characteristic of Graves’ disease is the infiltration of immune cells into the thyroid (Dayan et al., 1991; McIver and Morris, 1998; Totterman et al., 1979). Overall, this analysis suggests a role for SDV expression in sex-biased diseases.

### Genes with sex-differentially variable expression show evidence of increased selective constraint acting on gene expression and sequence

Given that genes with sex-biased expression (SBE) have been shown to be associated with relaxed selective constraint at both the sequence and expression levels, we asked whether SDV genes would similarly be associated with reduced constraint (Gershoni and Pietrokovski, 2017; Naqvi et al., 2019). Stabilizing selection acting on gene expression is thought to reduce *cis*-regulatory variation and reduce the minor allele frequency (MAF) of *cis*-regulatory variants (Glassberg et al., 2019; Josephs et al., 2015). Therefore, if SDV expression is associated with reduced selective constraint, one would expect an enrichment of *cis*-regulatory variation in SDV genes. To determine whether SDV genes were enriched or depleted for *cis*-regulatory variation, we obtained expression quantitative trait loci (eQTL) summary data for each tissue from GTEx (v8). For each tissue with at least 20 autosomal SDV genes, we tested for enrichment of genes with significant eQTLs (eGenes, q-value<0.1) within that tissue’s set of SDV genes. Only genes that were tested in both studies in a given tissue were considered. Contrary to expectation, we observed a depletion of eGenes within the set of SDV genes in most tissues, and this depletion was significant in three tissues: sun-exposed skin, breast tissue, and pituitary glands (**Figure 3A**). We also tested whether the median fold enrichment of eGenes across the 20 tested tissues (median fold enrichment = 0.93) was lower than expected by chance. To this end, we randomly permuted the SDV labels in each tissue (genes were either SDV or not SDV in each tissue) and calculated a median fold enrichment across all tissues. In total 10,000 permutations were performed and for each round of permutation, the median fold enrichment of eGenes in the set of SDV genes was recorded. We found that the observed median fold enrichment of eGenes among SDV genes was significantly lower than expected by chance suggesting that expression levels of SDV genes are evolving under elevated constraint (p=1.3e-03, permutation test) (**Figure 3B).**

**Figure 3.**
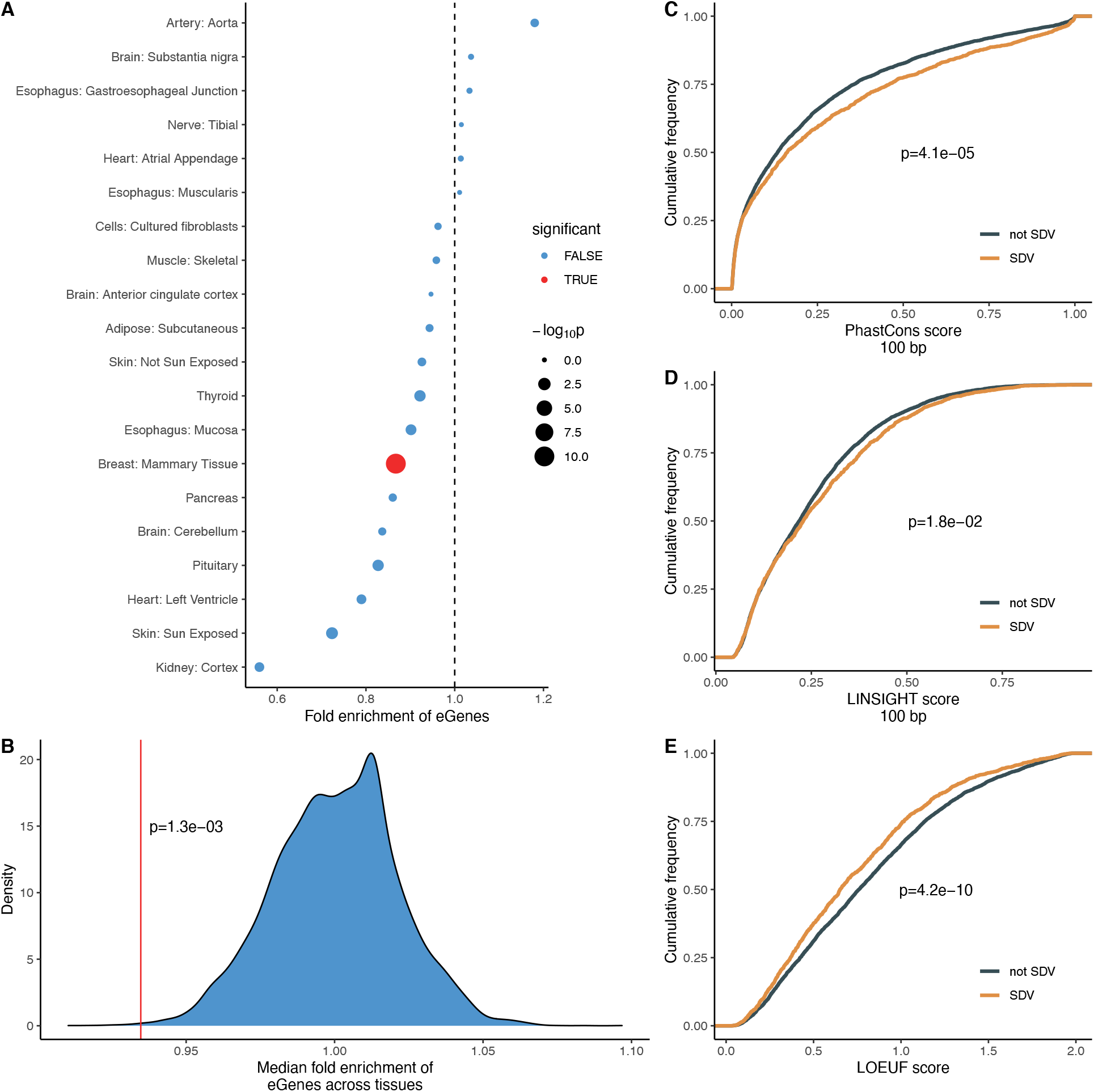
SDV genes are associated with increased constraint. A) 14 out of 20 tested tissues with at least 20 SDV genes show a depletion in eGenes among SDV genes. Significance was tested using Fisher’s exact test at FDR<0.05. B) Across the 20 tested tissues the median fold enrichment of eGenes is significantly lower than expected by chance. The red line shows the observed median fold-enrichment, while the blue distribution shows the empirical null distribution. C) Genes with SDV expression in at least one tissue show significantly elevated mean phastCons scores in proximal promoters (100 bp upstream of TSS) indicating increased conservation. D) Genes with SDV expression in at least one tissue show significantly increased LINSIGHT scores, indicating greater functional importance. E) Genes with SDV expression in at least one tissue show significantly reduced LOEUF scores, indicating more intolerance to loss of function mutations. For C-E, SDV genes were considered to be SDV if they showed SDV expression in at least one tissue out of the 20 tested tissues. Only genes expressed in all 20 tested tissues were considered to remove bias. For C-E, p-values were calculated using the Mann-Whitney U test.

We examined if the observed depletion of *cis*-regulatory variation in SDV genes in breast tissue could be explained by their elevated cell-type specificity. For example, one could imagine that most expression variation (across bulk samples) in cell-type specific genes is driven by sample-to-sample differences in cell-type composition and not by *cis*-regulatory genetic factors. Therefore, one might expect to observe a depletion of eQTLs in cell-type specific genes. Upon examination, we found no significant relationship between cell-type specificity and the presence or absence of a significant eQTL (p=0.99, Mann-Whitney *U* test)(**Supplementary Figure 7**).

Although our analysis of eQTL data provided some evidence that expression levels of SDV genes are evolving under increased constraint, a depletion of *cis*-regulatory variation might also be expected if the gene expression variation observed in SDV genes was due to environmental factors and/or *trans*-regulatory genetic variation to a larger extent than in non-SDV genes. We therefore sought to quantify selective constraint using a complementary—and more direct—approach. To this end, we used 20-species phastCons scores to look for further evidence that the *cis*-regulatory regions of SDV genes were evolutionary highly constrained (Siepel et al., 2005). In particular, we examined the sequence conservation in the proximal promoter regions of genes –100 bp upstream from the transcription start sites (TSSs). The phastCons score of a nucleotide can be interpreted as the probability that the nucleotide resides in a conserved element (Siepel et al., 2005). We found that genes with SDV expression in at least one tissue showed significantly elevated phastCons scores compared to genes which did not exhibit SDV expression in any tissues (p=4.1e-05, Mann-Whitney U test) (**Figure 3C**). Only genes expressed in all tissues with at least 20 SDV genes were considered to remove bias, i.e., a gene is more likely to be SDV if it’s expressed in more tissues and genes expressed in many tissues tend to be more evolutionarily constrained (Karczewski et al., 2020).

Similarly, we also examined the LINSIGHT scores of SDV genes. LINSIGHT scores combine information from intraspecific genetic polymorphism, interspecific genetic divergence, and functional genomic data to provide a probability that a mutation at a given base-pair will have deleterious fitness consequences (Huang et al., 2017). We found that genes with SDV expression in at least one tissue showed significantly higher mean LINSIGHT scores within their proximal promoter regions than genes without SDV expression (p=1.8e-02, Mann-Whitney U test).

Finally, we asked whether the increased constraint acting on the expression levels of SDV genes would be reflected in other metrics of dosage sensitivity. To do this, we compared LOEUF scores in SDV genes and non SDV genes. LOEUF (loss-of-function observed/expected upper bound fraction) is a score that reflects the tolerance of a protein to loss-of-function mutations (Karczewski et al., 2020). The scores range from 0 to 2, with higher scores indicating higher tolerance to loss-of-function mutations and therefore reduced selective constraint. We only considered genes expressed in all tissues with at least 20 SDV genes and found that genes with SDV in at least one tissue had significantly reduced LOEUF scores compared to genes without SDV expression in any tissue (p=4.2e-10, Mann-Whitney U test) (**Figure 3E**), corroborating that SDV genes tend to be more constrained at expression level.

### SDV genes in breast tissue show evidence of sex-specific genetic regulation

To determine if SDV genes showed evidence of sex-specific genetic regulation, we downloaded sex-biased eQTL (sbQTL) data from GTEx (Oliva et al., 2020). We restricted our analysis to breast tissue given that it was found to have both the largest number of SDV genes (in our study) and by far the largest number of sbQTLs of any tissue (Oliva et al., 2020). To test if genes with sbQTLs were enriched for SDV genes we performed a gene set enrichment analysis (GSEA). By only considering genes tested in both analyses and ranking genes according to their corrected −log_10_(p-values) in the sbQTL analysis we found that SDV genes were significantly enriched at lower p-values (p=1.2e-2, permutation test) (**Supplementary Figure 8**). Additionally, we found that all 13 SDV genes with higher variation in males and significant sbQTLs (FDR<0.25, the threshold used in Oliva et al., 2020) had eQTLs with larger effect sizes in males than females. This represents a significant enrichment of genes with larger eQTL effect sizes in males relative to the set of all genes with significant sbQTLs (p=7.4e-4, hypergeometric test). These results indicate that SDV genes are enriched for sex-specific genetic architectures regulating gene expression and suggest that SDV expression may have an underlying genetic component.

## Discussion

In this work, we examined a largely unappreciated aspect of transcriptomic sex differences: genes with sex differences in inter-individual expression variation. In particular, we focused on genes that showed sex differences in inter-individual expression overdispersion. We termed these genes sex-differentially variable (SDV) genes. Across 43 tested tissues, we found that roughly 18% of expressed genes showed SDV patterns of expression in at least one tissue (**Figure 1B**). Our results were undoubtedly affected by the unequal number of samples in different tissues – we had more power to detect SDV genes in some tissues than others. Nevertheless, we identified more SDV genes in breast tissue than in any other tissue. Issues of sample size aside, from a physiological perspective this seems logical since the breast is the only tissue tested known to have functional significance in only one sex: the production of milk is essentially female-specific. Other studies have found evidence that low inter-individual transcriptional variation in a gene is indicative of increased constraint acting on the gene (Fair et al., 2020). Since the vast majority (94%) of SDV genes identified in breast tissue showed higher expression variability in males than in females, this is consistent with the idea that expression levels should be more functionally constrained in female breast tissue than in male breast tissue.

To gain biological insight into the nature of SDV genes we performed a gene set overrepresentation analysis followed by meta-analysis. We found that SDV genes were enriched for genes from sets related to sex hormone response, immune response, and various signaling and cancer-associated pathways (**Figure 2A, B**). Additionally, we found that SDV genes were enriched for the targets of three transcription factors implicated in cancer (**Figure 2C**). The overrepresentation of SDV genes in gene sets related to various biological pathways and transcription factors indicates that SDV expression may be caused by SDV activity at the pathway level. In breast tissue, the overrepresentation of genes with high variability in males in sets related to estrogen response suggests that genes of particular biological importance to one sex may show higher variation in the opposite sex.

In order to further examine the biological significance of SDV genes in breast tissue we analyzed previously published single-cell RNA sequencing data of breast epithelial tissue. We found that SDV genes (identified in the GTEx data) were expressed in a more cell-type specific manner (**Figure 2D**). These results indicate that SDV genes may be more likely to be involved in cell-type specific functions. Additionally, this suggests that sex differences in gene expression variability may be partially driven by sex differences in cellular composition variability across individuals.

To examine a possible link between SDV expression and disease, we analyzed summary GWAS data from a classically female-biased disease: Graves’ disease. We found a significant enrichment of Graves’ associated genes in the set of SDV genes with higher variability in females in healthy thyroid tissue (**Figure 2E**). In the future, the role of SDV expression in sex-biased diseases should be thoroughly investigated by examining expression in both healthy and diseased tissue.

We found that SDV genes tended to be depleted for *cis*-regulatory variation overall (**Figure 3A. B**). Additionally, we showed that SDV genes show significantly increased constraint in both proximal promoter and protein-coding sequences. These results are consistent with increased constraint acting on the expression levels of SDV genes and suggest that SDV genes may be evolving under increased sex-specific constraint acting on gene expression.

We propose that SDV expression can arise through sex-specific increases in stabilizing selection acting on gene expression. To test this, we performed a forward Monte Carlo simulation of the evolution of a gene evolving under sex-specific stabilizing selection (**Figure 4**). Our model assumes that the expression depends on the additive contributions of a shared genetic locus (this locus contributes equally to both sexes), two sex-specific loci (each locus only contributes to expression in one sex), and noise (to model environmental effects) (see methods). Therefore, an assumption of the model is some degree of sex-specific genetic architecture for SDV genes, which is supported by empirical evidence (**Supplementary Figure 8**). The three loci were in linkage equilibrium. Increased stabilizing selection in one sex results in a difference in inter-individual phenotypic variances between the two sexes – with the sex experiencing increased selection having relatively reduced variation (**Figure 4A, B**). Additionally, increased selection in one sex results in reduced genetic variation in the set of shared genetic loci – consistent with our observations of reduced *cis*-regulatory variation in SDV genes (**Figure 4C**). These results are qualitatively consistent with predictions from quantitative genetics theory. Within a single generation, stronger stabilizing selection in one sex – for example, females – would lead to reduced additive genetic variance in females compared to males. Assuming that phenotypic variance is the sum of additive genetic and environmental variances, and that environmental variances are the same in both sexes, it implies that phenotypic variance will be lower in females than in males. Additionally, increased stabilizing selection in females will decrease the additive genetic covariance between the sexes to a larger extent than identical stabilizing selection in both sexes. This corresponds to our observation of reduction in the variance of the shared genetic locus under conditions of increased selection in females (**Figure 4C**). A more detailed description of the theoretical predictions is presented in the **Appendix**.

**Figure 4.**
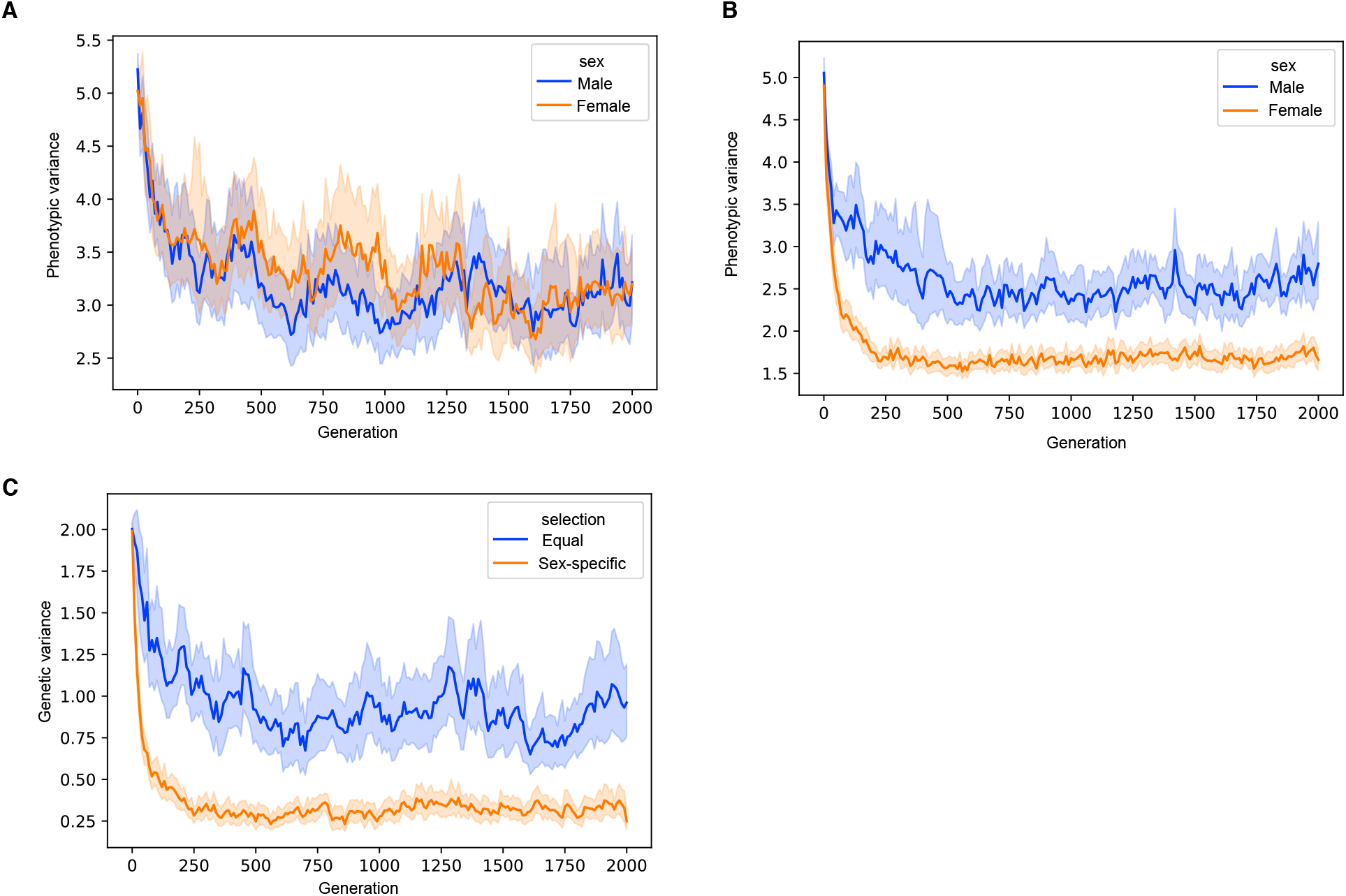
Simulations showing that SDV expression can arise through sex-specific constraint. Simulation results showing the output from 30 simulations in both the equal selection and sex-specific selection conditions. A) Equal constraint in both sexes results in similar phenotypic variance in both sexes. B) Increased sex-specific constraint acting in females reduces the phenotypic (expression) variation across individuals. C) Increased constraint in females results in reduced genetic variance, consistent with our observations of reduced *cis*-regulatory variation in SDV genes.

Although sex-specific selection resulting in decreased expression variance in the sex with increased selection is one possible explanation for SDV expression, it is not the only possibility. Alternatively, SDV expression, in particular genes with higher variability in females, could be a result of hormonal fluctuations as part of the menstrual cycle. The overrepresentation of estrogen response genes among female SDV genes supports this as a possible explanation (**Figure 2A**). These genes may also evolve under increased selective constraint in females. Both explanations for SDV expression suggest sex-specific or sex-biased functions for SDV genes, which could help explain the observation that a majority of SDV genes also exhibit SBE expression in a few tissues (**Supplementary Figure 4**).

Overall, our results suggest that genes with sex differences in inter-individual gene expression variability are possibly involved in sex-specific biological functions and may be evolving under sex-specific constraint acting on gene expression. These results highlight the possible importance of SDV genes in a functional and evolutionary sense, but future work needs to be done to properly understand the role of SDV genes in human health.

## Methods

### Power Simulations

Random samples were generated from negative binomial distributions using the ‘R’ function ‘rnegbin’ from the library ‘MASS’. To simulate the effect of ‘sex’ on the overdispersion parameter, samples were drawn from two negative binomial distributions with equal mean (*μ*) parameters and different overdispersion (*σ*, or equivalently 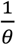 in the ‘MASS’ documentation) parameters. Similarly, to determine false positive rates samples were drawn from identical distributions. To restrict the search space, *μ* was fixed at 1500 for all simulations and the size of both samples combined was fixed at 400. These parameters were typical of the gene-expression levels and sample sizes found in GTEx data. Moreover, varying the mean parameter only marginally affected power calculations. For the unbalanced group sizes simulations, the size of one group was 320 while the size of the other was 80. Both the baseline overdispersion parameter (*σ*) and the difference in overdispersion between groups (Δ*σ*) were varied over the course of the simulation experiments. The variances of the two samples were compared using the Fligner-Killeen test (‘fk.test’ function from the ‘R’ library ‘npsm’), Levene’s test (‘levene.test’ function from the ‘R’ library ‘lawstat’), and a likelihood-ratio test implemented using negative binomial GAMLSS models. The more complex GAMLSS model parametrized the mean (*μ*) and the overdispersion (*σ*) parameters as functions of sex, *σ* ~ *sex* and *μ* ~ *sex*, while the simplified model parametrized only the mean in terms of sex with the overdispersion being modeled as constant. The p-value for the significance of the sex overdispersion term was saved. At each parameter combination 1000 sample pairs were drawn and p-values from each of the three tests were saved. Power was calculated as the fraction of 1000 simulations where the p-value fell below an alpha of 0.05. False positive rate was calculated as the faction of 1000 simulations where the p-value fell below and alpha of 0.05 when the two underlying distributions being sampled for a sample pair were identical. It was observed that power for Levene’s test markedly improved when log-transforming the sampled count data, while the Fligner-Killeen test showed largely unaltered power: hence samples were transformed as ln(counts + 1) before testing with Levene’s test and the Fligner-Killeen test. Raw samples were used for the GAMLSS likelihood-ratio test.

### Identification of SDV genes

For each tissue, SDV genes were identified separately using RNA-seq count data downloaded from GTEx-v8 (dbGAP accession number # phs000424.v8.p2). To reduce confounding due to population structure, only European Americans were considered. First, counts per million (CPM) values were calculated using edgeR and genes with no expression (CPM=0) in at least one individual were removed. To identify individuals that were global expression outliers, dimensionality reduction was then performed on log_2_ transformed CPM values using principal component analysis (PCA) with the ‘R’ function ‘prcomp’. Since outliers can greatly affect estimates of variation, we removed individuals whose global expression profiles deviated significantly from other individuals in a given tissue. Individuals whose global expression patterns were more than 1.5 interquartile ranges higher than the third quartile or lower than the first quartile along any principal component explaining more than 5% of the total variance in the dataset were considered outliers and removed from further analysis.

Following global outlier removal three negative binomial models were fit to the count data (one gene at a time) using gamlss. The first and most complex model (model 1) parametrized the mean and non-Poisson variation terms of the negative binomial distribution (*μ* and *σ*, respectively) as *μ ~ sex* + *age* + *ischemic time* + *RIN* + *PC1* + *PC2* + *PC3* + *PC4* + *PCS* + *offset*(*offset*) and *σ ~ sex* + *age* + *ischemic time* + *RIN* + *PC1* + *PC2* + *PC3* + *PC4* + *PCS*, where *RIN* represents RNA integrity number and the 5 PCs represent genotype principal components obtained from GTEx. The offset term, to account for library size was calculated as *log*(*library size* * *normalization factors*), where the normalization factors were calculated using the ‘calcNormFactors’ function from edgeR using the ‘upperquartile’ method. Model 2 had the same form as model 1 for *μ*, however for the formula for the non-Poisson variation was *σ ~ age*+*ischemic time* + *RIN* + *SPCs*. Model 3 had the same formula as model 1 for *σ* however the formula for the mean was *μ ~ age*+*ischemic time* + *RIN* + *SPCs* + *offset*(*offset*). To test for the significance of the effect of sex on non-Poisson variation (*p_σ,sex_*) a likelihood ratio test was performed comparing model 1 and model 2. To test for the significance of the effect of sex on mean expression (*p_μ,sex_*) a likelihood ratio test was performed comparing model 1 and model 3. The coefficients corresponding to the sex terms for both *μ* and *σ*(*β_μ,sex_* and *β_σ, sex_*) from model 1 were recorded for each gene. After every gene was tested in a given tissue p-values for *μ* and *σ* were corrected independently using the Benjamini-Hochberg (BH) procedure using the ‘R’ function ‘p.adjust’.

Unsurprisingly we noticed that individual expression outliers at the gene level dramatically affected our estimates of *p_σ,sex_* and *β_σ,sex_*. To account for this and to generally validate the robustness of our models we performed randomization tests for all genes with a BH corrected *p_σ,sex_* <0.05. For each gene the sex labels were randomly permuted, model 1 was fit to the randomized data, and *β_σ,sex_* values were recorded. We performed 1000 permutations for every gene to generate an empirical null distribution for *β_σ,sex_*. An empirical p-value was calculated as the fraction of random permutations where *abs*(*β_σ,sex,perm_*) > *abs*(*β_σ,sex,obs_*). These empirical p-values were then corrected using the BH procedure. In the end genes were considered SDV if the original BH corrected p-values (*p_σ,sex_*) were less than 0.05 and the BH corrected empirical p-values were less than 0.05.

### Hallmark enrichment analysis

Gene set annotations for collection ‘H’ (hallmark gene sets) were downloaded from MSigDB using the ‘R’ function ‘msigdbr’ from the package ‘msigdbr’. Only tissues with at least 20 male or female high variability genes were considered. Enrichment of gene sets for male high variability and female high variability genes was calculated separately. Enrichment p-values for each gene set in each tissue were calculated using a hypergeometric test (using the ‘R’ function ‘phyper’). For each gene set a meta-analysis was performed across all tested tissues using a variation of Fisher’s method. This approach was implemented as follows. For each gene set a test statistic was calculated by taking −2 times the sum of the log p-values in each tested tissue. In an ordinary implementation of Fisher’s method this test statistic would have a chi-squared distribution under the null hypothesis. However, given that the underlying p-values are based on a discrete test (the hypergeometric test) a core assumption of Fisher’s method is violated: namely that for an individual tissue p-values should be uniformly distributed under the null hypothesis. To overcome this, we generated empirical null distributions for the test statistic described above using a permutation approach: within each tissue we permuted the labels of each gene (not SDV, female high variability, and male high variability) and calculated a p-value for the label of interest (female high variability or male high variability) using a hypergeometric test. This permutation procedure was done for every tissue tested – finally resulting in a null value for the sum of log p-values test statistic described above. For each gene set 10,000 such permutations were performed to generate an empirical null distribution for the test statistic. The empirical p-values for each gene set were calculated by finding the fraction of permutations resulting in a test statistic larger than the one observed. These combined p-values across gene sets were corrected using the BH procedure and gene sets with corrected p-values<0.05 were considered significant. Gene sets that were highly tissue/developmentally specific were not tested in the meta-analysis. The 7 removed hallmark gene sets were pancreas beta cells, adipogenesis, myogenesis, spermatogenesis, bile acid metabolism, heme metabolism, and xenobiotic metabolism.

### Transcription factor enrichment analysis

The analysis was performed in a fashion identical to the hallmark enrichment analysis except that the gene sets from collection ‘C3’ subcollection ‘GTRD’ were used.

### Single-cell RNA sequencing analysis

The count matrix for individual 4 was downloaded from GSE113197 (Nguyen et al., 2018). All clustering and marker identification analysis was performed using the ‘R’ library ‘Seurat’ version 3.2.3. Cells which had fewer than 500 and more than 6000 expressed features were removed. Cells with more than 10% of their transcripts coming from mitochondrial genes were removed. The counts were normalized using the function ‘SCTransform’. Both dimensionality reduction (using the function ‘RunUMAP’) and clustering (function: ‘FindClusters’) were performed using the first 10 principal components of the normalized counts. L1, L2, and basal cells were identified based on their expression of markers “KRT18”, “SPLI”, and “KRT14”. Clusters expressing “KRT18” and “SPLI” were deemed to be L1 cells, clusters only expressing “KRT18” were deemed to be L2 cells, and clusters expressing “KRT14” were taken to be basal cells.

To calculate the cell-type specificity of each gene raw counts were aggregated within each of the three cell types, generated 3 pseudo-bulk count vectors (one for each cell type). For each cell-type CPMs were calculated by multiplying the pseudo-bulk count vector by 1 million and dividing by the sum of the vector. The CPM values were then log_2_ transformed. Cell-type specificity was calculated using the standard metric for tissue specificity 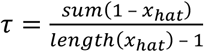 where *x_hat_* is the vector representing a given gene’s expression across the 3 cell-types divided by the maximum expression of that gene in any of the three cell-types.

### Graves’ disease analysis

Summary GWAS statistics were obtained from the NHGRI-EBI GWAS catalog for study EFO_0004237, downloaded on 08/04/2022 (Buniello et al., 2019).

### eQTL analysis

Summary statistics for single-tissue eQTL mapping performed in European-American individuals were downloaded from the GTEx-v8 website. Genes with at least one eQTL with q-value<0.1 were considered to be eGenes. A two-sided Fisher’s exact test, as implemented in the ‘fisher.test’ function, was used to test for the enrichment/depletion of eGenes among SDV genes in each tissue. Only tissues with at least 20 autosomal SDV genes were considered. Only autosomal genes with expression high enough to be tested in both the original eQTL study and our study were considered. To test if the median fold enrichment of eGenes across tissues was significant we randomly permuted the SDV labels in each tissue and calculated a median fold enrichment across all tissues. In total 10,000 permutations were performed and for each round of permutation the median fold enrichment of eGenes across the 20 tissues was recorded. The empirical p-value was calculated as the fraction of all permutations where the fold enrichment was at least as low as the observed fold enrichment.

### PhastCons analysis

Genome-wide phastCons scores (in hg38 coordinates) generated from a 20 species alignment were downloaded from http://hgdownload.cse.ucsc.edu/goldenPath/hg38/phastCons20way/. Transcription start sites (TSS) for all annotated genes were downloaded from BioMart using the ‘R’ package biomaRt (Durinck et al., 2005, 2009). The most distal TSS was used for each gene. Only genes expressed in all tissues containing at least 20 SDV genes were tested (20 tissues overall). A gene was considered ‘SDV’ if it displayed ‘SDV’ expression in at least 1 of the 20 tested tissues, otherwise it was considered to be ‘not SDV’. Requiring genes to be expressed in all 20 tissues was done to remove bias, i.e., a gene is more likely to be SDV if it’s expressed in more tissues and genes expressed in many tissues tend to be more evolutionarily constrained (Karczewski et al., 2020). Further increasing the number of tissues (by decreasing the minimum number of SDV genes allowed in each tissue) reduces power by decreasing the intersection of genes expressed across all tissues. Mean phastCons scores were obtained for each gene in a region 100 bp upstream from their TSSs.

### LINSIGHT analysis

Data was downloaded from the Siepel Laboratory website: https://siepellab.labsites.cshl.edu/software/. The data was processed and analyzed in the same manner as the phastCons data following conversion to hg38 coordinates.

### Loss of function analysis

Loss of function metrics were downloaded from https://gnomad.broadinstitute.org/downloads. In particular the v2.1.1 loss of function metrics by gene were used. The analysis was similar to the analysis for PhastCons score: only genes expressed across all tissues with at least 20 SDV genes were considered. LOEUF scores were compared between ‘SDV’ and ‘not SDV’ genes.

### Sex-specific stabilizing selection simulation

The simulation assumed a diploid, diecious, and randomly mating population of 500 individuals. In each individual, the phenotype (expression) was modelled as the additive contribution of 3 genetic loci and environmental noise. One of the genetic loci had equal effect in both sexes while each of the two other genetic loci only had effect in one sex or the other. The allelic effects of each genetic locus in each individual were initialized as draws from a standard normal distribution (mean=0 and standard deviation=1). The environmental component was also taken to be a random number drawn from a standard normal distribution. Similarly, mutation resulted in the addition of a number drawn from a standard normal distribution to the allelic effect and the mutation rate was set to 5e-3 per generation at each genetic locus in each individual. The fitness of each individual was taken to be *w* = *e*^−*s*(*G*+*E*)^2^^, where ‘*s*’ represented the strength of selection, *‘G’* was the total sum of genetic allelic effects (from the shared and sex-specific locus), and ‘*E*’ was the environmental effect. For the simulations with equal selection in both sexes ‘*s’* was 0.005, while in simulation with sex-specific selection ‘*s’* was 0.005 in males and 0.05 in females. Selection only affected viability and had no impact on fecundity. An individual survived to reproduce if a random draw from a uniform distribution on [0, 1] was less than or equal to the individual’s fitness; therefore, the fitness represents the probability of an individual surviving to reproduce. In each generation 100 males and 100 females were chosen and paired at random with replacement. Each pair produced 5 offspring and the sex of each offspring was determined randomly. The simulations were run for a total of 2000 generations. Every 10 generations the variance of the phenotype (across individuals in the population) was recorded along with the variance of the allelic effect of the shared genetic component. For both the equal selection and sex-specific selection models 30 independent simulations were run.

### GSEA analysis of sbQTL data

Sex-biased QTL (sbQTL) data from breast tissue was downloaded from GTEx v8. Genes were only considered if they were tested in our analysis and in the sbQTL analysis. Genes were ranked according to corrected −log_10_(p-values) where the (Bonferroni) correction was made to account for multiple SNPs tested per gene. GSEA analysis was performed using the ‘R’ function ‘fgsea’ from the library ‘fgsea’(Korotkevich et al., 2021).

## Appendix

### Theory behind sex-specific stabilizing selection

The evolution of a single quantitative trait in males and females can be predicted from the 2-by-2 additive genetic variance-covariance matrix:

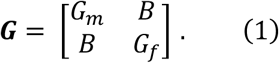

Where the diagonal terms represent the additive genetic variances in males and females, while the identical off-diagonal terms represent the additive genetic covariance between the sexes (Barker et al., 2010; Lande, 1980). In the absence of directional selection, the change in the genetic variance-covariance matrix due to stabilizing selection over the course of a generation is (Barker et al., 2010; Phillips and Arnold, 1989):

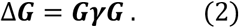

Where *γ* is a matrix describing the strength of stabilizing selection within each sex:

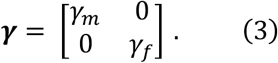

For stabilizing selection to occur the diagonal terms must be negative with decreasing values indicating stronger stabilizing selection. Substituting (3) and (1) into (2) yields:

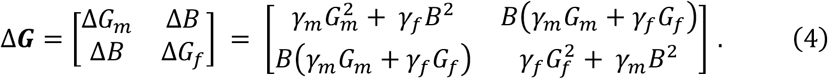

By assuming that initially the trait has identical genetic variance in males and females (which is assumed at the start of our simulation): we can use the simplification *G_m_* = *G_f_* = *G*. This leads to the following equations for Δ*G_m_* and Δ*G_f_*:

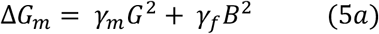

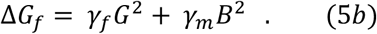

In the case of stronger stabilizing selection in females: *γ_f_* < *γ_m_* < 0. Additionally, theoretical and empirical evidence suggests that generally the magnitude of intra-sex genetic variance is larger than the magnitude of inter-sex genetic covariance: *G* > *B* (Barker et al., 2010). Therefore, (5*a, b*) yields: Δ*G_f_* < Δ*G_m_* < 0. Following the assumptions made in our simulation (namely that the phenotypic variance in each sex is just the sum of the environmental variances and the additive genetic variances (*G*), and that the environmental variance is the same in both sexes) leads to the result that phenotypic variance will decrease to a larger extent in females than in males over the course of a generation. This is what we observed (however, over multiple generations) in our simulation (**Figure 4A, B**).

To further relate our simulation results to the theory described above we can formulate the phenotypes and their variances from the simulation as follows:

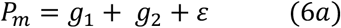

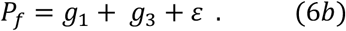

*P_m_* and *P_f_* are the phenotypes in males and females respectively. *g*_1_, *g*_2_, and *g*_3_ are the genetic contributions of the shared genetic locus, the male specific genetic locus, and the female specific genetic locus. The environmental contribution to the phenotypes is *ε*. From (6*a, b*), since all the phenotypic components are independent (so their covariances are 0), the phenotypic variances are:

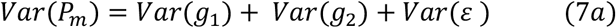

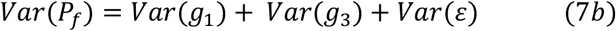

and the phenotypic covariances are:

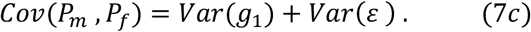

From (7*a, b, c*) it becomes apparent that *G_m_* corresponds to *var*(*g*_1_) + *var*(*g*_2_), *G_f_* = corresponds to *var*(*g*_1_) + *var*(*g*_3_), and *B* corresponds to *var*(*g*_1_). From the off-diagonal terms in equation (4) it is evident that increasing the strength of selection in females (decreasing *γ_f_*) makes Δ*B* more negative – which is consistent with the observed relative reduction in the genetic variance in the shared locus (*var*(*g*_1_)) (**Figure 4C**).

## Data access and code availability

The code generated for this work has been deposited on GitHub: https://github.com/LiZhaoLab/SDV.

## Acknowledgments

We thank Zhao lab members for discussions about the project, Jason Mezey and Christina Leslie for helpful discussions during early stages of the project, Mike Love for statistical advice, and Andy Clark for the suggestion of power analysis. We thank Matt Hahn and Ziyue Gao for their comments and suggestions for a recent version of the manuscript. We thank Jason Banfelder and Bala Jayaraman from The Rockefeller University High Performance Computing (HPC) Center for their dedicated work in these extraordinary times.

## References

Avery, J.T., Zhang, R., and Boohaker, R.J. (2021). GLI1: A Therapeutic Target for Cancer. Front. Oncol. 11.

Barker, B.S., Phillips, P.C., and Arnold, S.J. (2010). A test of the conjecture that G-matrices are more stable than B-matrices. Evolution 64, 2601–2613.

Beery, A.K., and Zucker, I. (2011). Sex bias in neuroscience and biomedical research. Neurosci. Biobehav. Rev. 35, 565–572.

Buniello, A., MacArthur, J.A.L., Cerezo, M., Harris, L.W., Hayhurst, J., Malangone, C., McMahon, A., Morales, J., Mountjoy, E., Sollis, E., et al. (2019). The NHGRI-EBI GWAS Catalog of published genome-wide association studies, targeted arrays and summary statistics 2019. Nucleic Acids Res. 47, D1005–D1012.

Catalán, A., Hutter, S., and Parsch, J. (2012). Population and sex differences in Drosophila melanogaster brain gene expression. BMC Genomics 13, 654.

Chella Krishnan, K., Mehrabian, M., and Lusis, A.J. (2018). Sex differences in metabolism and cardiometabolic disorders. Curr. Opin. Lipidol. 29, 404–410.

Choleris, E., Galea, L.A.M., Sohrabji, F., and Frick, K.M. (2018). Sex differences in the brain: Implications for behavioral and biomedical research. Neurosci. Biobehav. Rev. 85, 126–145.

Chu, X., Pan, C.-M., Zhao, S.-X., Liang, J., Gao, G.-Q., Zhang, X.-M., Yuan, G.-Y., Li, C.-G., Xue, L.-Q., Shen, M., et al. (2011). A genome-wide association study identifies two new risk loci for Graves’ disease. Nat. Genet. 43, 897–901.

Clement, V., Sanchez, P., de Tribolet, N., Radovanovic, I., and Ruiz i Altaba, A. (2007). HEDGEHOG-GLI1 Signaling Regulates Human Glioma Growth, Cancer Stem Cell Self-Renewal, and Tumorigenicity. Curr. Biol. 17, 165–172.

Cooper, J.D., Simmonds, M.J., Walker, N.M., Burren, O., Brand, O.J., Guo, H., Wallace, C., Stevens, H., Coleman, G., Franklyn, J.A., et al. (2012). Seven newly identified loci for autoimmune thyroid disease. Hum. Mol. Genet. 21, 5202–5208.

Dayan, C.M., Londei, M., Corcoran, A.E., Grubeck-Loebenstein, B., James, R.F., Rapoport, B., and Feldmann, M. (1991). Autoantigen recognition by thyroid-infiltrating T cells in Graves disease. Proc. Natl. Acad. Sci. 88, 7415–7419.

Durinck, S., Moreau, Y., Kasprzyk, A., Davis, S., De Moor, B., Brazma, A., and Huber, W. (2005). BioMart and Bioconductor: a powerful link between biological databases and microarray data analysis. Bioinformatics 21, 3439–3440.

Durinck, S., Spellman, P.T., Birney, E., and Huber, W. (2009). Mapping identifiers for the integration of genomic datasets with the R/Bioconductor package biomaRt. Nat. Protoc. 4, 1184–1191.

Fair, B.J., Blake, L.E., Sarkar, A., Pavlovic, B.J., Cuevas, C., and Gilad, Y. (2020). Gene expression variability in human and chimpanzee populations share common determinants. Elife 9, e59929.

Falconer, D.S. (1967). The inheritance of liability to diseases with variable age of onset, with particular reference to diabetes mellitus. Ann. Hum. Genet. 31, 1–20.

Gershoni, M., and Pietrokovski, S. (2017). The landscape of sex-differential transcriptome and its consequent selection in human adults. BMC Biol. 15, 7.

Glassberg, E.C., Gao, Z., Harpak, A., Lan, X., and Pritchard, J.K. (2019). Evidence for Weak Selective Constraint on Human Gene Expression. Genetics 211, 757LP–772.

Hao, Y., Hao, S., Andersen-Nissen, E., Mauck, W.M., Zheng, S., Butler, A., Lee, M.J., Wilk, A.J., Darby, C., Zagar, M., et al. (2020). Integrated analysis of multimodal single-cell data. BioRxiv 2020.10.12.335331.

Huang, Y.-F., Gulko, B., and Siepel, A. (2017). Fast, scalable prediction of deleterious noncoding variants from functional and population genomic data. Nat. Genet. 49, 618–624.

Itoh, Y., and Arnold, A.P. (2015). Are females more variable than males in gene expression? Meta-analysis of microarray datasets. Biol. Sex Differ. 6, 1–9.

de Jong, T. V, Moshkin, Y.M., and Guryev, V. (2019). Gene expression variability: the other dimension in transcriptome analysis. Physiol. Genomics 51, 145–158.

Josephs, E.B., Lee, Y.W., Stinchcombe, J.R., and Wright, S.I. (2015). Association mapping reveals the role of purifying selection in the maintenance of genomic variation in gene expression. Proc. Natl. Acad. Sci. 112, 15390 LP–15395.

Karczewski, K.J., Francioli, L.C., Tiao, G., Cummings, B.B., Alföldi, J., Wang, Q., Collins, R.L., Laricchia, K.M., Ganna, A., Birnbaum, D.P., et al. (2020). The mutational constraint spectrum quantified from variation in 141,456 humans. Nature 581, 434–443.

Khodursky, S., Svetec, N., Durkin, S.M., and Zhao, L. (2020). The evolution of sex-biased gene expression in the Drosophila brain. Genome Res. 30, 874–884.

Klein, S.L., and Flanagan, K.L. (2016). Sex differences in immune responses. Nat. Rev. Immunol. 16, 626–638.

Kolmykov, S., Yevshin, I., Kulyashov, M., Sharipov, R., Kondrakhin, Y., Makeev, V.J., Kulakovskiy, I. V, Kel, A., and Kolpakov, F. (2021). GTRD: an integrated view of transcription regulation. Nucleic Acids Res. 49, D104–D111.

Korotkevich, G., Sukhov, V., Budin, N., Shpak, B., Artyomov, M.N., and Sergushichev, A. (2021). Fast gene set enrichment analysis. BioRxiv 60012.

Lande, R. (1980). Sexual Dimorphism, Sexual Selection, and Adaptation in Polygenic Characters. Evolution (N. Y). 34, 292.

Liberzon, A., Birger, C., Thorvaldsdóttir, H., Ghandi, M., Mesirov, J.P., and Tamayo, P. (2015). The Molecular Signatures Database (MSigDB) hallmark gene set collection. Cell Syst. 1, 417–425.

Lopes-Ramos, C.M., Chen, C.-Y., Kuijjer, M.L., Paulson, J.N., Sonawane, A.R., Fagny, M., Platig, J., Glass, K., Quackenbush, J., and DeMeo, D.L. (2020). Sex Differences in Gene Expression and Regulatory Networks across 29 Human Tissues. Cell Rep. 31, 107795.

Mauvais-Jarvis, F., Bairey Merz, N., Barnes, P.J., Brinton, R.D., Carrero, J.-J., DeMeo, D.L., De Vries, G.J., Epperson, C.N., Govindan, R., Klein, S.L., et al. (2020). Sex and gender: modifiers of health, disease, and medicine. Lancet 396, 565–582.

McCarthy, D.J., Chen, Y., and Smyth, G.K. (2012). Differential expression analysis of multifactor RNA-Seq experiments with respect to biological variation. Nucleic Acids Res. 40, 4288–4297.

McIver, B., and Morris, J.C. (1998). The pathogenesis of graves’ disease. Endocrinol. Metab. Clin. North Am. 27, 73–89.

Meisel, R.P., Malone, J.H., and Clark, A.G. (2012). Faster-X Evolution of Gene Expression in Drosophila. PLOS Genet. 8, e1003013.

Nakabayashi, K., Tajima, A., Yamamoto, K., Takahashi, A., Hata, K., Takashima, Y., Koyanagi, M., Nakaoka, H., Akamizu, T., Ishikawa, N., et al. (2011). Identification of independent risk loci for Graves’ disease within the MHC in the Japanese population. J. Hum. Genet. 56, 772–778.

Naqvi, S., Godfrey, A.K., Hughes, J.F., Goodheart, M.L., Mitchell, R.N., and Page, D.C. (2019). Conservation, acquisition, and functional impact of sex-biased gene expression in mammals. Science. 365, eaaw7317.

Nguyen, Q.H., Pervolarakis, N., Blake, K., Ma, D., Davis, R.T., James, N., Phung, A.T., Willey, E., Kumar, R., Jabart, E., et al. (2018). Profiling human breast epithelial cells using single cell RNA sequencing identifies cell diversity. Nat. Commun. 9, 2028.

Ober, C., Loisel, D.A., and Gilad, Y. (2008). Sex-specific genetic architecture of human disease. Nat. Rev. Genet. 9, 911–922.

Oliva, M., Muñoz-Aguirre, M., Kim-Hellmuth, S., Wucher, V., Gewirtz, A.D.H., Cotter, D.J., Parsana, P., Kasela, S., Balliu, B., Viñuela, A., et al. (2020). The impact of sex on gene expression across human tissues. Science. 369, eaba3066.

Parisi, M., Nuttall, R., Naiman, D., Bouffard, G., Malley, J., Andrews, J., Eastman, S., and Oliver, B. (2003). Paucity of genes on the Drosophila X chromosome showing male-biased expression. Science 299, 697–700.

Parsch, J., and Ellegren, H. (2013). The evolutionary causes and consequences of sex-biased gene expression. Nat. Rev. Genet. 14, 83–87.

Phillips, P.C., and Arnold, S.J. (1989). Visualizing Multivariate Selection. Evolution (N. Y). 43, 1209–1222.

Quaranta, R., Pelullo, M., Zema, S., Nardozza, F., Checquolo, S., Lauer, D.M., Bufalieri, F., Palermo, R., Felli, M.P., Vacca, A., et al. (2017). Maml1 acts cooperatively with Gli proteins to regulate sonic hedgehog signaling pathway. Cell Death Dis. 8, e2942–e2942.

Ranz, J.M., Castillo-Davis, C.I., Meiklejohn, C.D., and Hartl, D.L. (2003). Sex-dependent gene expression and evolution of the Drosophila transcriptome. Science (80-.). 300, 1742–1745.

Rao, R.C., and Dou, Y. (2015). Hijacked in cancer: the KMT2 (MLL) family of methyltransferases. Nat. Rev. Cancer 15, 334–346.

Rigby, R.A., and Stasinopoulos, D.M. (2005). Generalized additive models for location, scale and shape. J. R. Stat. Soc. Ser. C (Applied Stat. 54, 507–554.

Robinson, M.D., and Smyth, G.K. (2007). Moderated statistical tests for assessing differences in tag abundance. Bioinformatics 23, 2881–2887.

Robinson, M.D., McCarthy, D.J., and Smyth, G.K. (2010). edgeR: a Bioconductor package for differential expression analysis of digital gene expression data. Bioinformatics 26, 139–140.

Sakaue, S., Kanai, M., Tanigawa, Y., Karjalainen, J., Kurki, M., Koshiba, S., Narita, A., Konuma, T., Yamamoto, K., Akiyama, M., et al. (2021). A cross-population atlas of genetic associations for 220 human phenotypes. Nat. Genet. 53, 1415–1424.

Siepel, A., Bejerano, G., Pedersen, J.S., Hinrichs, A.S., Hou, M., Rosenbloom, K., Clawson, H., Spieth, J., Hillier, L.W., Richards, S., et al. (2005). Evolutionarily conserved elements in vertebrate, insect, worm, and yeast genomes. Genome Res. 15, 1034–1050.

Simonovsky, E., Schuster, R., and Yeger-Lotem, E. (2019). Large-scale analysis of human gene expression variability associates highly variable drug targets with lower drug effectiveness and safety. Bioinformatics 35, 3028–3037.

Soldin, O.P., and Mattison, D.R. (2009). Sex differences in pharmacokinetics and pharmacodynamics. Clin. Pharmacokinet. 48, 143–157.

Subramanian, A., Tamayo, P., Mootha, V.K., Mukherjee, S., Ebert, B.L., Gillette, M.A., Paulovich, A., Pomeroy, S.L., Golub, T.R., Lander, E.S., et al. (2005). Gene set enrichment analysis: A knowledge-based approach for interpreting genome-wide expression profiles. Proc. Natl. Acad. Sci. U. S. A. 102, 15545–15550.

Sun, Y., Wang, Y., Fan, C., Gao, P., Wang, X., Wei, G., and Wei, J. (2014). Estrogen promotes stemness and invasiveness of ER-positive breast cancer cells through Gli1 activation. Mol. Cancer 13, 137.

Totterman, T.H., Andersson, L.C., and Häyry, P. (1979). Evidence for thyroid antigen-reactive t lymphocytes infiltrating the thyroid gland in graves’disease. Clin. Endocrinol. (Oxf). 11, 59–68.

Williams, T.M., and Carroll, S.B. (2009). Genetic and molecular insights into the development and evolution of sexual dimorphism. Nat. Rev. Genet. 10, 797–804.

Wu, L., Aster, J.C., Blacklow, S.C., Lake, R., Artavanis-Tsakonas, S., and Griffin, J.D. (2000). MAML1, a human homologue of Drosophila mastermind, is a transcriptional co-activator for NOTCH receptors. Nat. Genet. 26, 484–489.

Zajitschek, S.R., Zajitschek, F., Bonduriansky, R., Brooks, R.C., Cornwell, W., Falster, D.S., Lagisz, M., Mason, J., Senior, A.M., Noble, D.W., et al. (2020). Sexual dimorphism in trait variability and its eco-evolutionary and statistical implications. Elife 9.

Zhao, S.-X., Xue, L.-Q., Liu, W., Gu, Z.-H., Pan, C.-M., Yang, S.-Y., Zhan, M., Wang, H.-N., Liang, J., Gao, G.-Q., et al. (2013). Robust evidence for five new Graves’ disease risk loci from a staged genome-wide association analysis. Hum. Mol. Genet. 22, 3347–3362.

