## Supplementary files for "Sex differences in inter-individual gene expression variability across human tissues"

### Supplementary figures for
### Sex differences in inter-individual gene expression variability across human tissues

**Authors:**

Samuel Khodursky[1], Caroline S. Jiang[2], Eric B. Zheng[1], Roger Vaughan[2], Daniel R. Schrider[3], Li Zhao[1,*]

**Affiliations:**

[1]Laboratory of Evolutionary Genetics and Genomics, The Rockefeller University, New York, NY 10065, USA
[2]Department of Biostatistics, The Rockefeller University, New York, NY 10065, USA
[3]Department of Genetics, University of North Carolina, Chapel Hill, NC 27514, USA

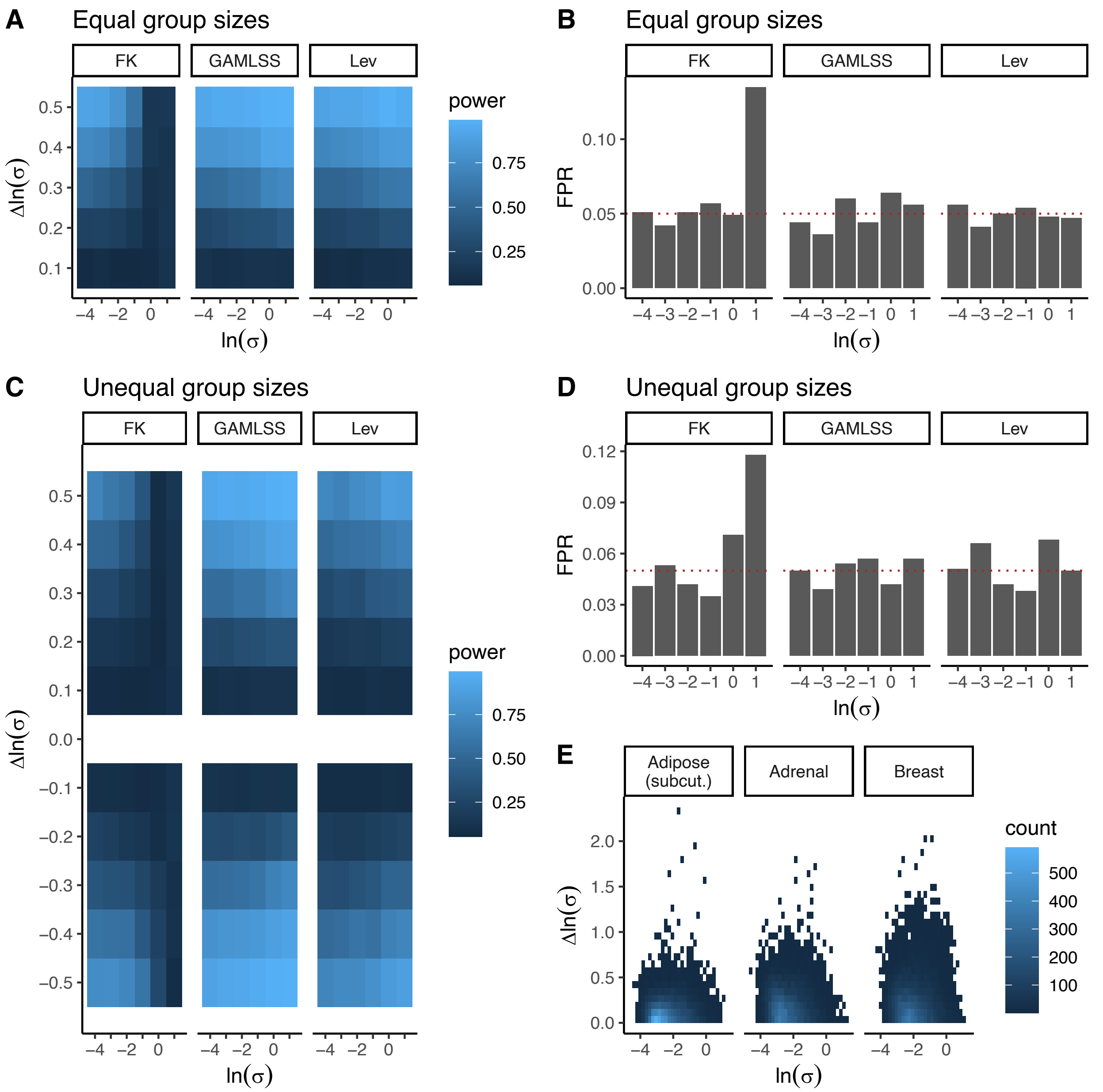


**Supplementary Figure 1. Power analysis.** A) Power at different baseline overdispersions and effect sizes using simulated data from groups of equal sample size (n=200 each). B) False positive rate for simulations with equal sample sizes. C) Power using simulated data from groups of unequal sample size (320 vs. 80). Positive effect sizes refer to an increase in variance in the larger group. D) False positive rate for simulations with unequal sample sizes. E) Observed overdispersion and effect sizes in data. For all analyses power was calculated at alpha=0.05.

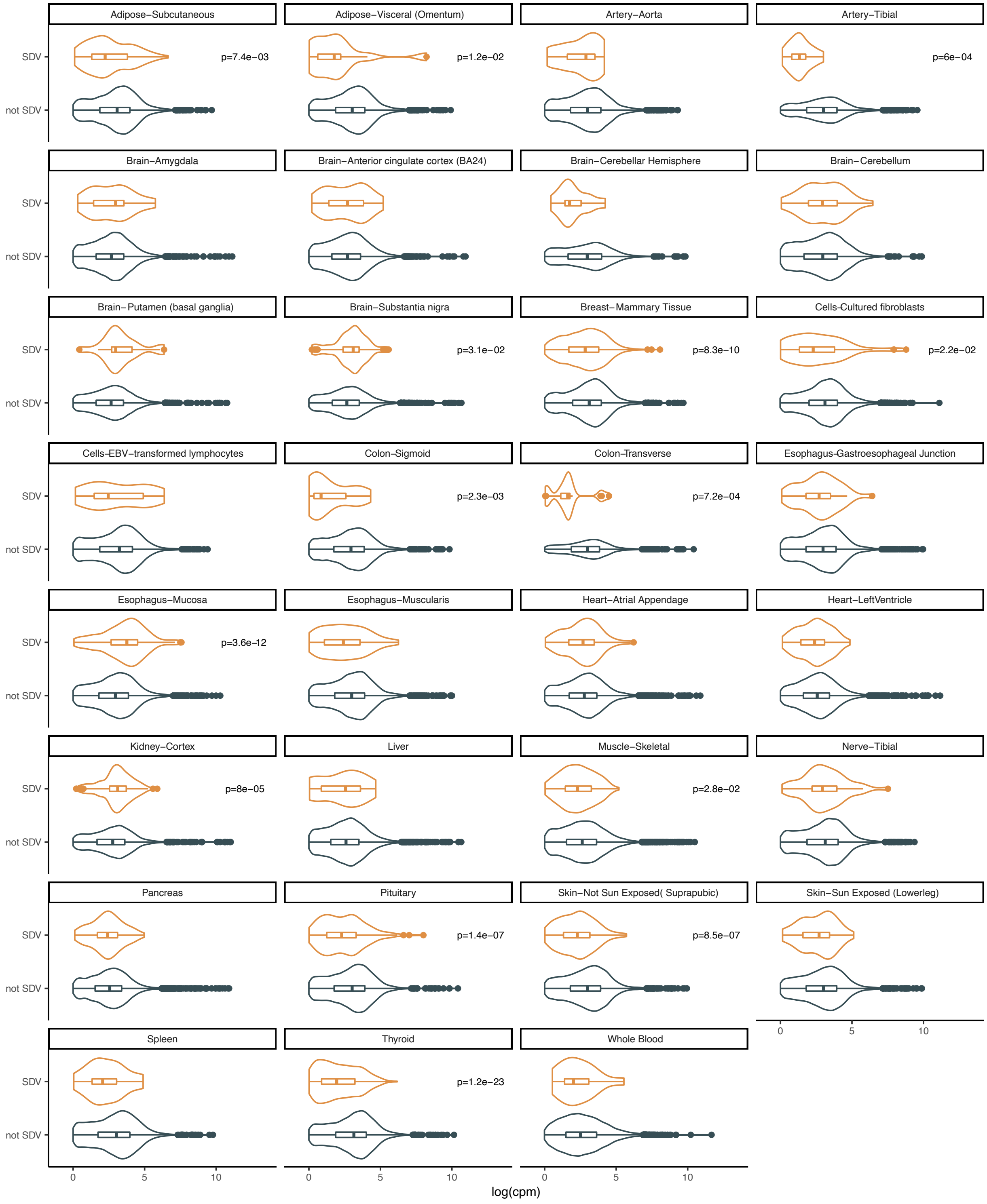


**Supplementary Figure 2A. Expression levels in SDV genes (CPM).** Significance tested using Mann-Whitney U test.

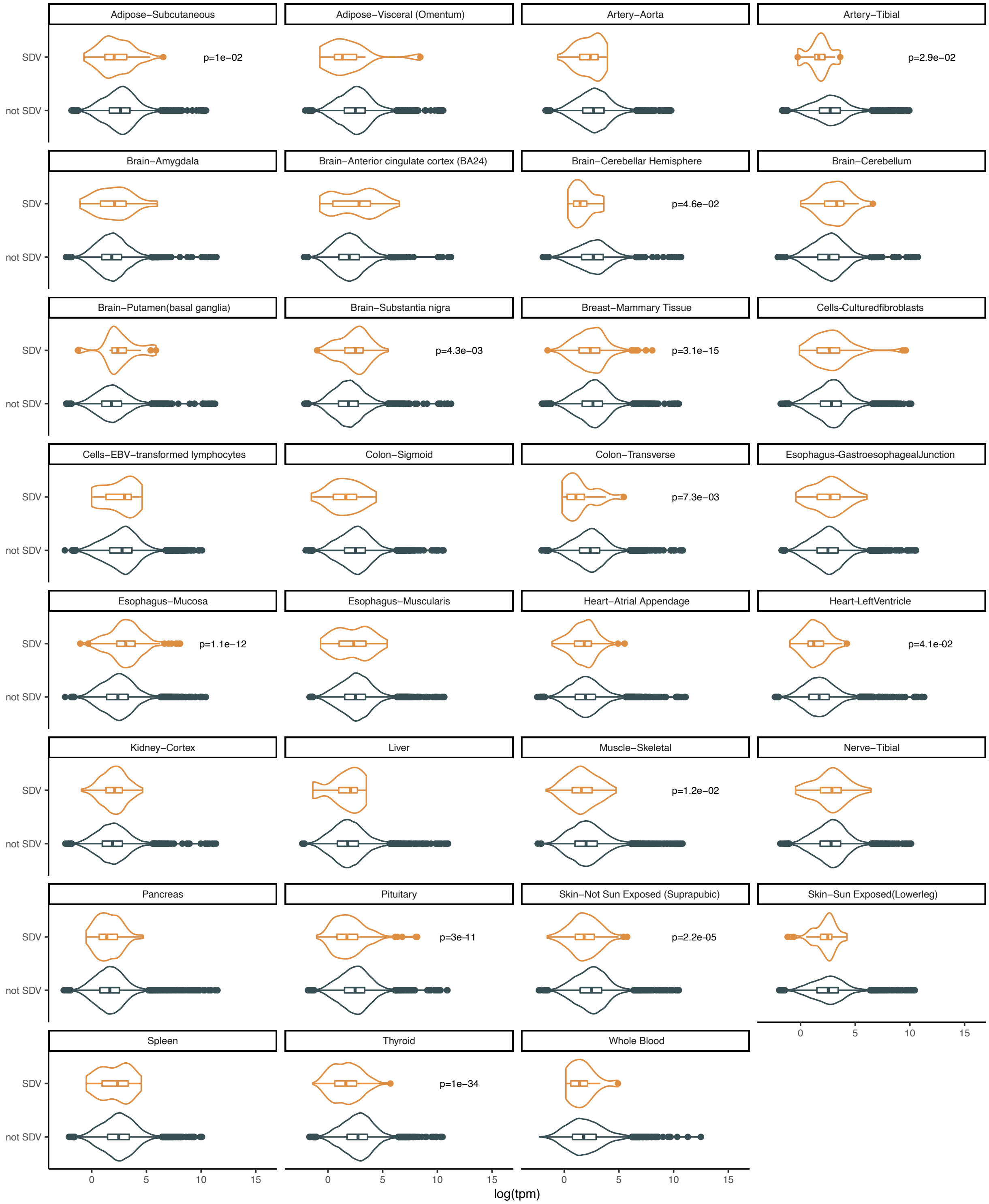


**Supplementary Figure 2B. Expression levels in SDV genes (TPM).** Significance tested using Mann-Whitney U test.

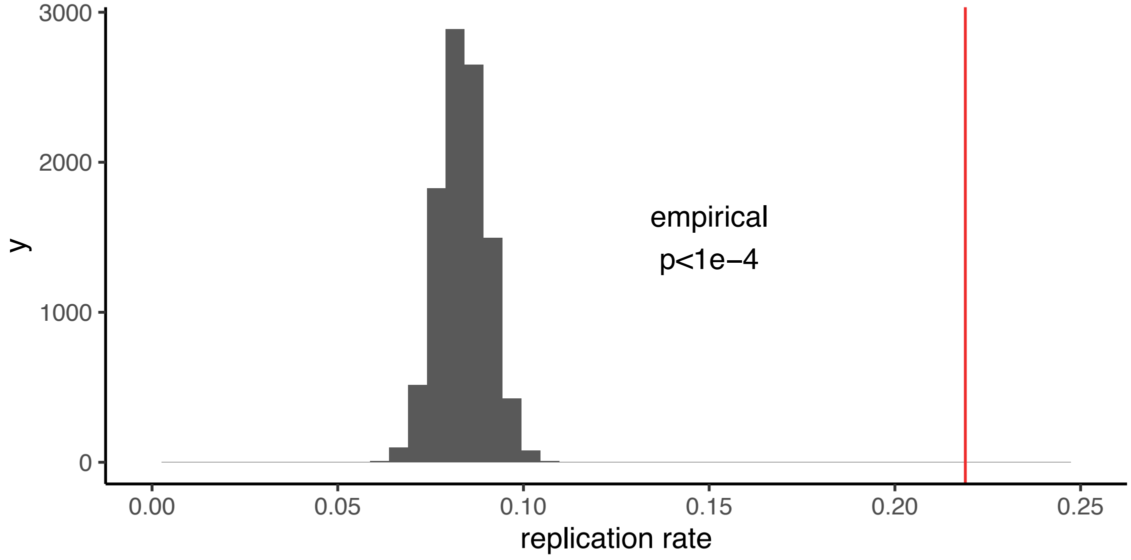


**Supplementary Figure 3. Replication of genes with significant sex differences in overdispersion in breast tissue.** The histogram shows the empirical null distribution for the replication rate while the red line shows the observed replication rate. Discovery was performed in a dataset containing 70% of the individuals and replication was performed in a dataset containing the remaining 30%.

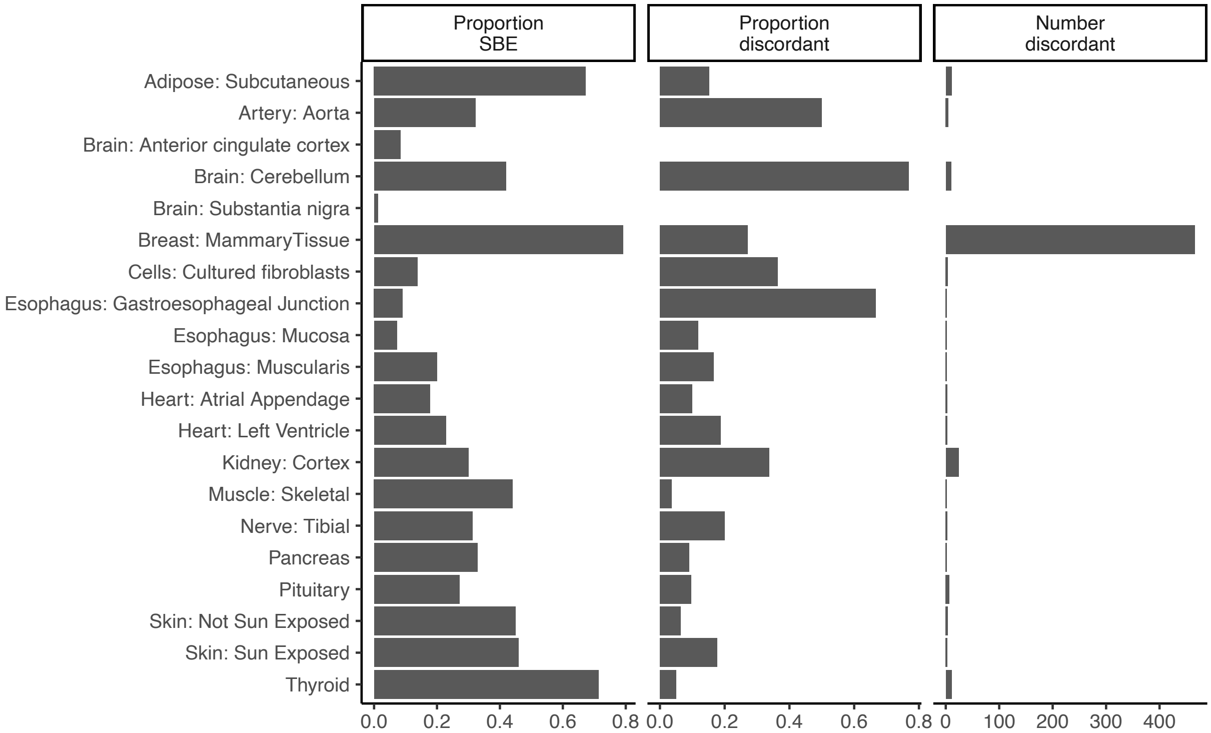


**Supplementary Figure 4. Relationship between genes with sex differences in overdispersion and genes with sex differences in mean expression levels.** The first panel shows the proportion of SDV genes in each tissue with significant sex-biased expression (FDR < 0.05). The second panel shows the proportion of genes with significant sex differences in mean and overdispersion that are discordant in their direction: e.g., genes with higher overdispersion in males while simultaneously having lower mean expression in males. The third panel gives the raw number of discordant genes in each tissue. Breast tissue has more than 400 genes with discordant sex differences in mean and overdispersion. Only tissues with at least 20 SDV genes are shown.

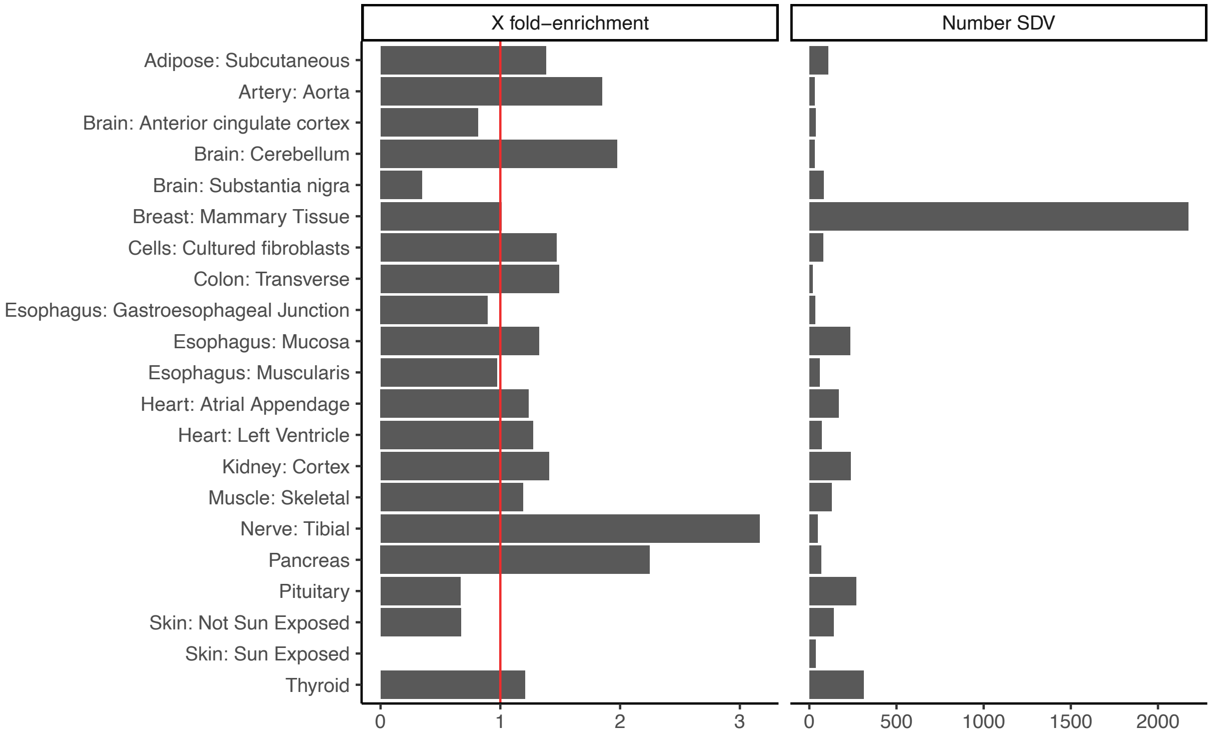


**Supplementary Figure 5. Enrichment of SDV genes on the X chromosome.**

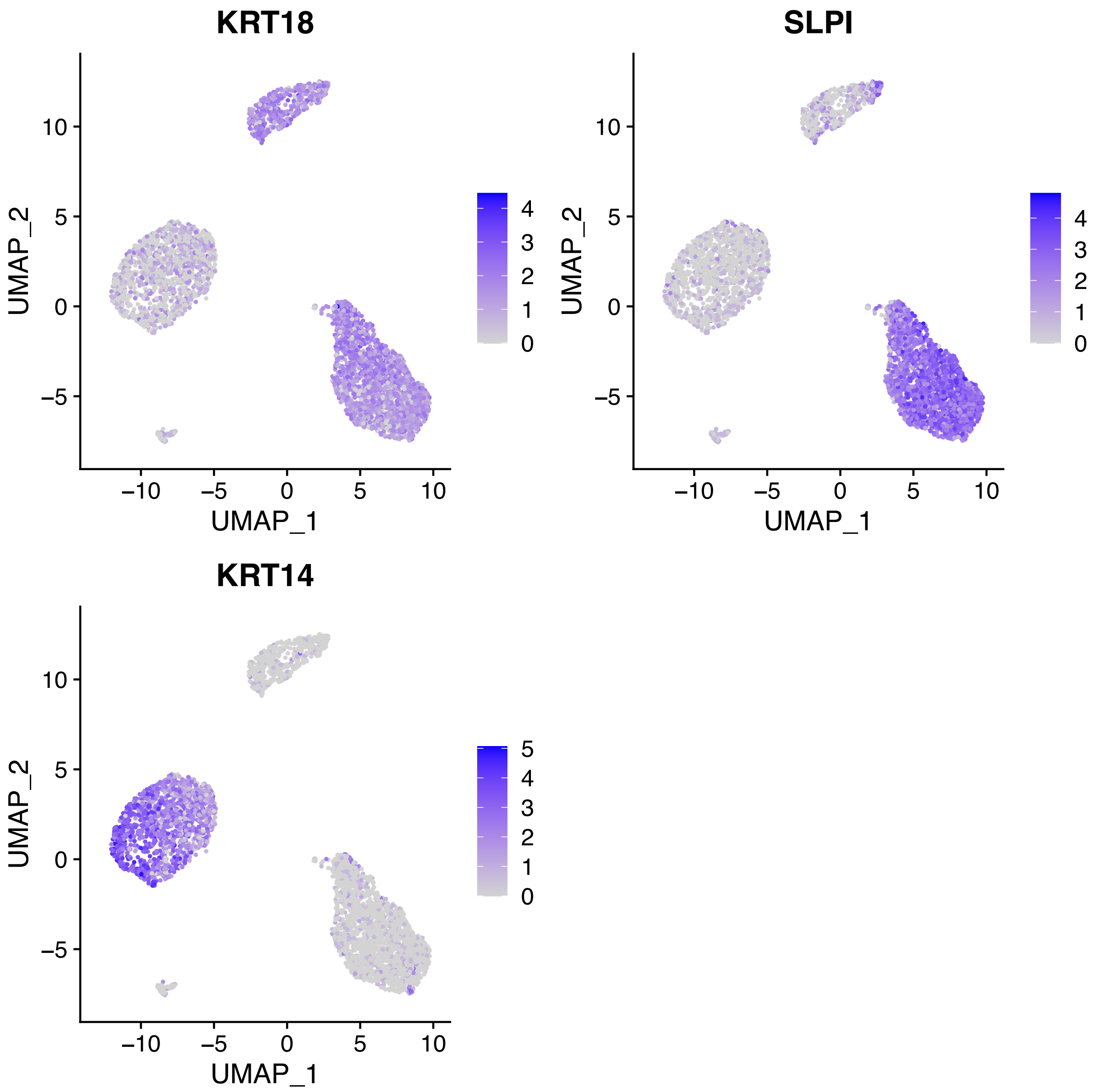


**Supplementary Figure 6. Dimensionality reduction of scRNA-seq data from individual 4 showing the 3 epithelial cell types.** UMAP projections show the three main cell types. The cluster expressing "KRT18" and "SPLI" was deemed to be L1 cells, the cluster only expressing "KRT18" was deemed to be L2 cells, and the cluster expressing "KRT14" was taken to be basal cells.

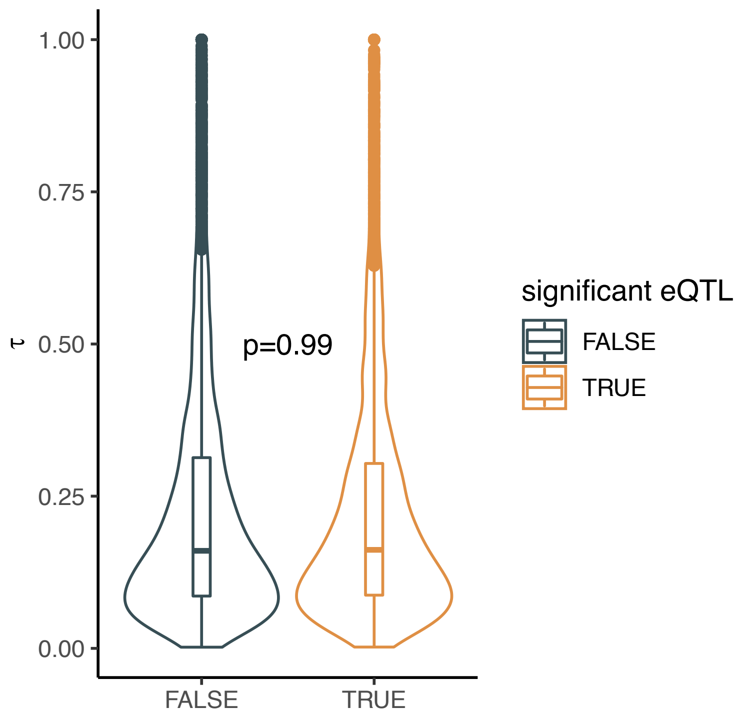


**Supplementary Figure 7. Relationship between eQTLs and cell-type specificity.** A) No relationship between cell-type specificity and the presence of eQTL in breast tissue**.** Significance tested using Mann-Whitney U test.

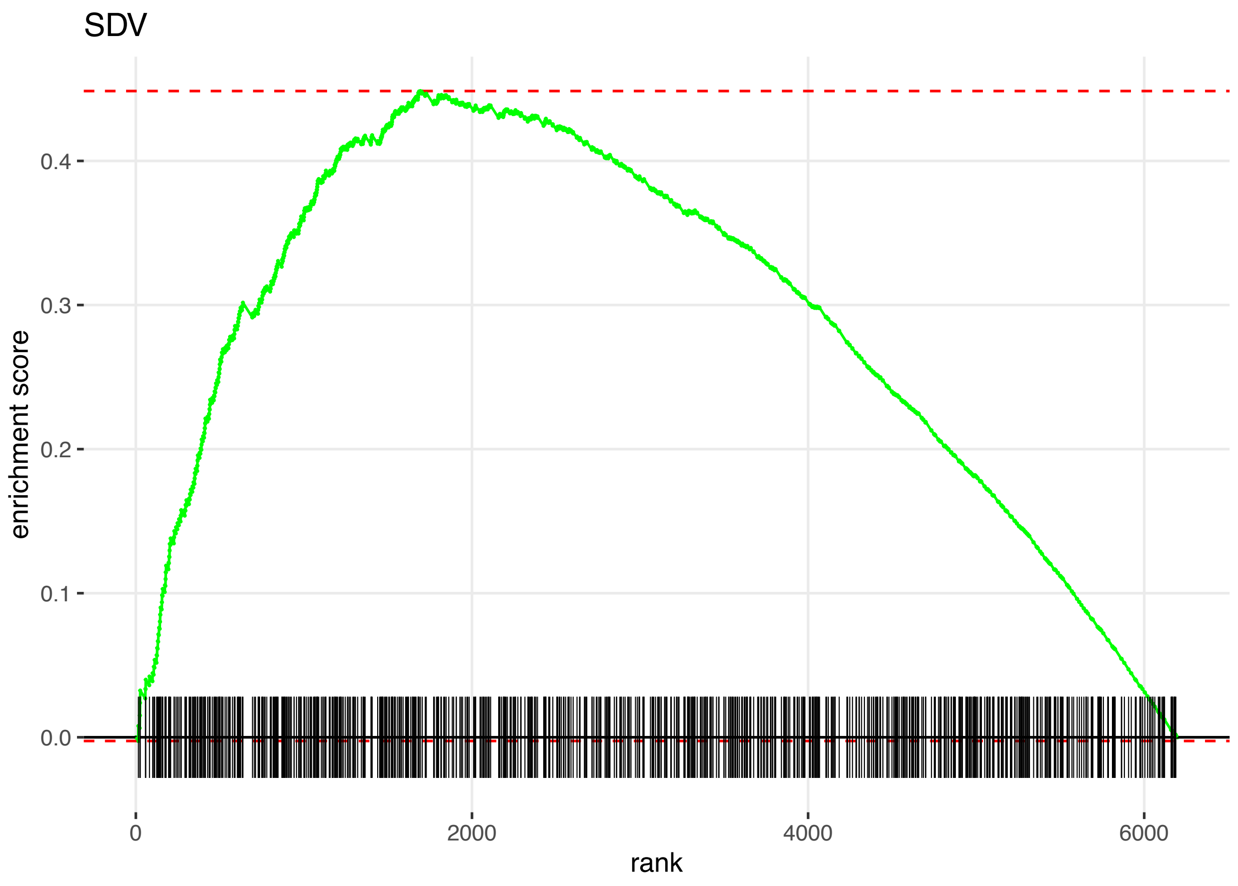


**Supplementary Figure 8. GSEA plot showing enrichment of SDV genes among genes with lower p-values in the sbQTL analysis of breast tissue.**
